# A generalizable speech neuroprosthesis

**DOI:** 10.64898/2026.07.23.739430

**Authors:** Zachery M. Fogg, Nicholas S. Card, Maitreyee Wairagkar, Aparna Srinivasan, Tyler Singer-Clark, Xianda Hou, Elizaveta Okorokova, Hamza Peracha, Carrina Iacobacci, Tiffany Brailow, Justin Jude, Hadar Levi-Aharoni, Trung Le, Domenick Mifsud, Pranav I. Deevi, Samuel R. Nason-Tomaszewski, Anna L. Pritchard, Yunnuo Zhang, Brice Richards, Payton Bechefsky, Leigh R. Hochberg, Ziv Williams, Kiarash Shahlaie, Nicholas AuYong, Daniel B. Rubin, Chethan Pandarinath, David M. Brandman, Sergey D. Stavisky

## Abstract

Intracortical brain-computer interfaces (BCIs) can restore communication to people with vocal tract paralysis by decoding cortical activity during attempted speech into text. State-of-the-art systems pairing neural-to-phoneme decoders with phoneme-to-word language models have achieved word error rates (WERs) as low as 1%, but only after collecting thousands of sentences of training data. Shortening the data collection process would facilitate scaling this new technology by reducing the time from device implant to high-accuracy communication. Here we introduce a transformer-based decoder model trained jointly across six intracortical speech BCI participants. For every participant — regardless of sex, disease etiology, or attempted speaking strategy — a multi-user model decoded speech more accurately (over 50% lower relative WER on average) than models trained on individual users’ data. Notably, the multi-user model could be finetuned on fewer than 200 sentences from a held-out user to achieve a WER below 7%. These results reveal how to pool intracortical data across people to yield more accurate, generalizable, and rapidly-deployable decoding models.

## 1. Introduction

Communication is central to quality of life and independence, but is very difficult for millions of people living with vocal-tract paralysis caused by neurological injury or disease, such as stroke or amyotrophic lateral sclerosis (ALS)^1–3^. Augmentative and assistive communication technologies, including eye-tracking and head-tracking systems, can partially restore communication but are typically slow, fatiguing, and unnatural ^4^. Brain-computer interfaces (BCIs)^5–8^, and speech BCIs^9–11^ in particular hold promise for fast, reliable, and more naturalistic communication for these populations.

Intracranial recordings from cortical regions involved in speech production can support speech BCIs that decode attempted speech directly from neural activity and produce communication output^12–25^. The most widely used “brain-to-text” paradigm to date has predominantly employed a neural phoneme decoder to translate intracortical signals from speech motor cortex into phonemes, which a language model then assembles into words and sentences displayed on a screen in real time^12–14,16,19–21^. Progress has been rapid: recent work demonstrated a speech BCI accurate and stable enough to support daily, independent, at-home communication in a clinical trial participant^13^.

Despite this progress, two obstacles limit the broad clinical deployment of speech BCIs. First, demonstrations of high-accuracy decoding have largely been confined to single participants or single disease etiologies, leaving generalizability of accurate decoding across more diverse patient populations underexplored. Second, achieving high accuracy — which we define here as approximately 10% word error rate (WER) or lower on a large vocabulary — has required first collecting thousands of sentences of participant-specific calibration data. Collecting these large datasets is effortful and time-consuming, especially for people with advanced neuromuscular disease^26^, and represents a major barrier to deploying speech BCIs as readily available assistive devices.

These obstacles motivate a central question: could training a single neural decoding model on data pooled from multiple intracortical speech BCI users both improve accuracy for existing users and reduce the calibration burden for new users? Evidence from adjacent fields suggests this is plausible. In automatic speech recognition, models such as Whisper^27^ and wav2vec 2.0^28^, trained on audio recordings from thousands of speakers, generalize to unseen users with little or no user-specific data. In surface electromyography (sEMG), a handwriting decoder trained on over 6,600 users generalized to new users without user-specific training^29^, and a typing decoder trained on over 100 users transferred to new users with minimal finetuning^30^. For *intracranial* speech BCIs, an early electrocorticography (ECoG) encoder–decoder reduced a new user’s data requirements by pretraining across users, but on closed sets of only a few dozen sentences^31^. Similarly, small-scale ECoG and stereoelectroencephalography (sEEG) studies on limited vocabularies found only modest cross-user gains^32,33^.

Whether cross-user transfer extends to action-potential-resolution neural measurements, however, is far from obvious. Intracortical recordings (which capture neuronal spiking activity in addition to coarser field potentials) are high-dimensional, differ substantially across individuals in electrode count and placement^34,35^, and lack a natural inter-subject alignment due to a near-random sampling of neurons’ functional properties^36^ (in contrast to the brain location-based alignment that bulk neural signals can exploit). Despite these regime-specific challenges, recent work suggests that brain activity shares a latent structure across individuals, and that, with enough data, deep learning models can extract it from single-unit recordings to support cross-subject transfer of non-speech motor decoders in mice^37^, non-human primates^38,39^, and humans^40^.

Cross-user transfer has been explored for *intracortical* speech BCIs only in restricted settings. Recent work found only modest benefits from transfer between users, and only in low-data regimes – gains that faded once more than 200 trials of a user’s own data were available^41^. A recent foundation-model approach went further, with multi-user pretraining improving offline decoding accuracy over single-user models for two users (a ∼20.0% relative WER improvement on attempted speech)^42^. However, these gains still required thousands of training sentences for each user, were shown only offline, and were demonstrated in two users – both with ALS. It thus remains unknown whether a single model trained on pooled intracortical datasets could decode large-vocabulary speech accurately across many diverse users, while reducing the calibration burden for new users.

We address this question by introducing a multi-user (MU) speech decoder trained simultaneously on neural data from six BrainGate2 clinical trial participants, two of whom are reported here for the first time. By learning shared phonemic neural representations across participants, the MU model achieves more accurate speech decoding than single-user (SU) baselines for every training participant, while remaining adaptable to new participants. In closed-loop use with three participants, MU models reduced WER by an average of 55.9% relative to SU models. In additional offline analyses across six participants, the MU model achieved a mean WER reduction of 51.1% relative to SU baselines, with one participant seeing a 73.3% reduction. The MU decoder’s accuracy also scaled with the number of pretraining participants, and the model adapted rapidly to held-out participants, achieving high-accuracy, large-vocabulary decoding from as little as 17 minutes of calibration data.

Together, these results suggest that the underlying latent state encoding of attempted speech in motor cortex is sufficiently conserved across individuals to support a shared decoding model, and establish multi-user training as a promising path toward generalizable BCIs that deliver high-accuracy communication with minimal calibration — moving the field closer to assistive devices that can be readily deployed to new users.

## 2. Results

### 2.1 Overview

This study includes data from six participants in the BrainGate2 clinical trial (ClinicalTrials.gov NCT00912041) with trial designators T12, T15, T16, T17, T21, and T22 (Table 1). All had dysarthria or anarthria due to ALS or brainstem stroke. Each participant had 1-6 microelectrode arrays (64 electrodes per array) surgically placed in the ventral precentral gyrus (speech motor cortex; Fig. S1). Arrays recorded intracortical activity via percutaneous wired connections. Additional participant details are provided in the Methods (Section 4.1).

**Table 1.**
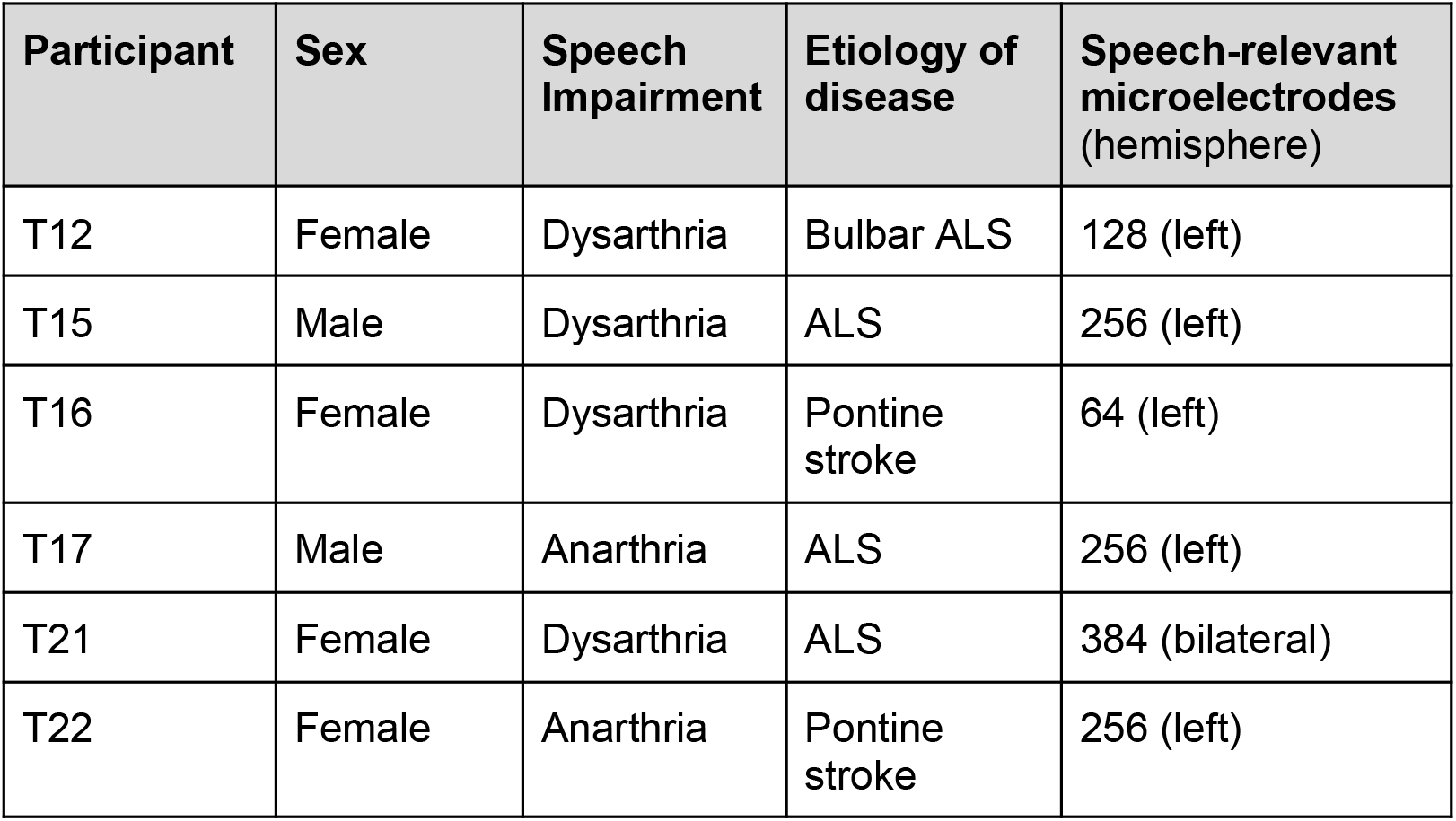
Participant information.

Voltage time-series were recorded and processed into neural features in real time as each participant attempted to speak. These features were decoded into the participant’s intended words with short latency (∼100-200 ms), displayed as text on a screen, and optionally played aloud at the end of each sentence via text-to-speech software. We collected speaking data from two tasks: instructed-delay Copy Tasks, in which the participant attempted to speak sentences displayed on a screen, and self-generated sentences without a prompt. As in previous work^12–14,16^, we used a cascaded decoder that first converted neural features into phoneme probabilities and then mapped phoneme sequences into words via a language model.

Our neural-to-phoneme decoder is a transformer-based architecture that can be trained either on data from a single user or jointly across multiple users (Fig. 1). During multi-user training, the model first maps each participant’s neural features through a learned, user-specific projection into a common latent space; from there, a single shared set of decoder weights processes every participant’s data identically, with no further user-specific processing. This shared space is a product of the training procedure used where every batch contains trials from all participants, forcing each gradient update to push the model toward a single, user-invariant solution.

**Figure 1.**
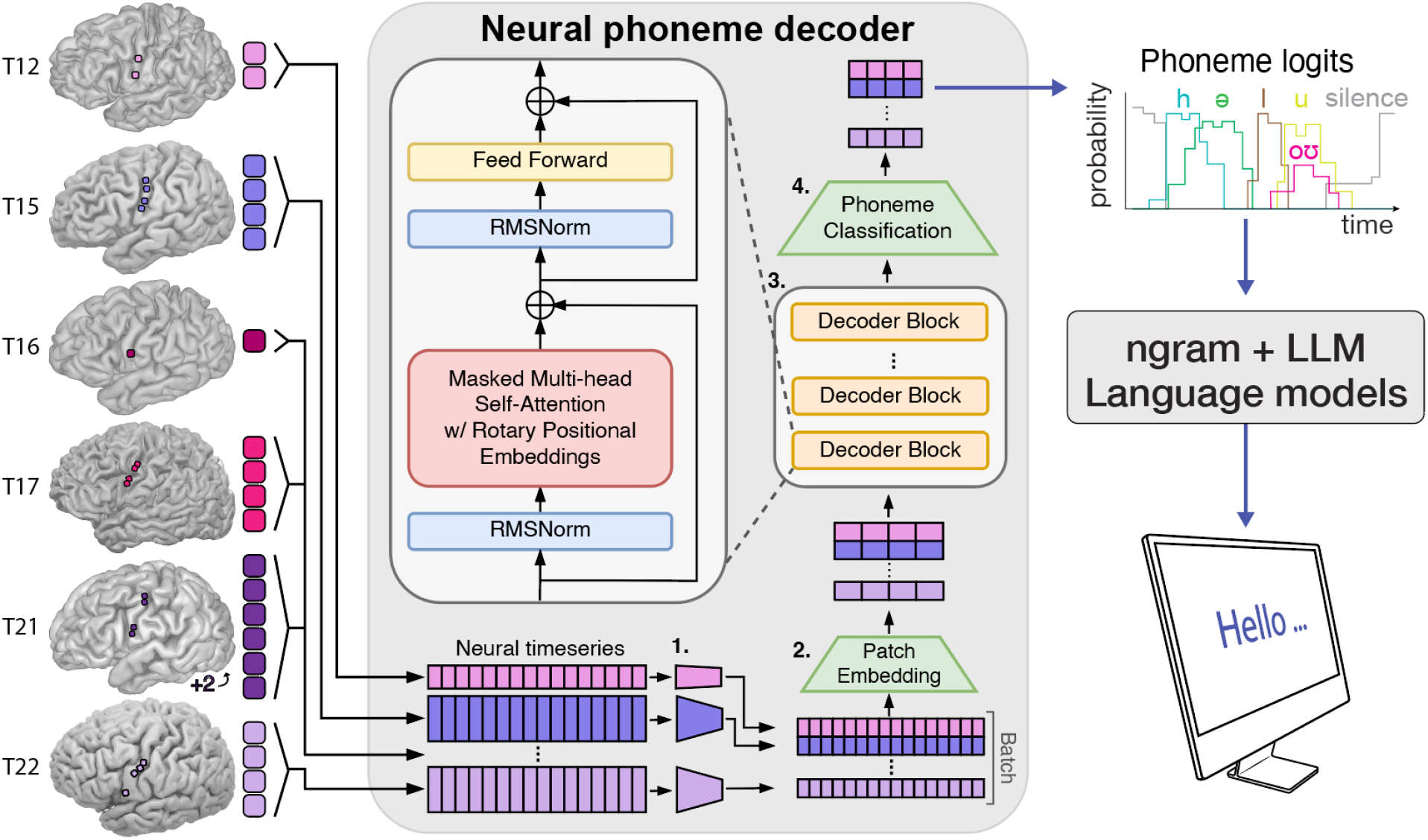
Overview of the multi-user brain-to-text speech BCI. Neural data is recorded from the speech motor cortex during attempted speech via 1 to 6 implanted microelectrode arrays in six participants. Neural data is processed into neural features (threshold crossings and spike-band power). A transformer decoder is pretrained on data from all six participants to predict spoken phoneme probabilities from neural features. During pretraining: (1) user-specific projections are learned to project each participant’s neural data into a shared latent space. (2) Each participant’s data is collated into a training batch and a patch embedding network downsamples the neural data from 20 ms to 80 ms resolution. (3) A stack of transformer decoder blocks processes features into rich, contextualized representations and then (4) phoneme probability distributions are predicted for each 80 ms bin. During real-time inference, a user’s neural data is transformed using their learned user-specific projection and decoded into phoneme probabilities that are continuously converted into the most likely words being spoken and displayed as text on a screen.

The neural phoneme decoder was pretrained on existing data, and could be continuously finetuned during closed-loop use for enhanced decoding stability, as in previous work^12,13^. For the phoneme-to-word language model, we used one of two implementations, each pairing an n-gram model with an LLM rescorer. The first is a previously-introduced weighted finite state transducer (WFST)^12–14^. The second is a new pipeline we built around Flashlight-Text^43^ and KenLM^44^, paired with an updated, finetuned LLM rescorer^45^; this implementation achieves similar decoding accuracy to the WFST pipeline, but is >3x faster and requires dramatically lower compute resources (Fig. S2). Full decoder architectures, hyperparameters, training, inference, and finetuning procedures are described in the Methods and Supplement.

### 2.2 Multi-user models improve closed-loop decoding

We performed closed-loop evaluation sessions with three participants (T16, T21, and T22) to compare decoding accuracy between SU and MU models (Figure 2; Table S1). In each session, participants attempted to speak prompted Switchboard sentences in a Copy Task (see Methods 4.3). Each closed-loop evaluation session used either a SU or a MU neural-to-phoneme decoder. SU models were trained on all available data from the target participant prior to the session; MU models were trained on the same within-participant data plus data from up to five other participants (see Methods 4.4.2). To enable direct comparison between the two model types, we also performed offline re-inference of the same sessions with both model types, using an offline simulation of continuous finetuning to match the online implementation (see Methods 4.5 and Table S2).

**Figure 2.**
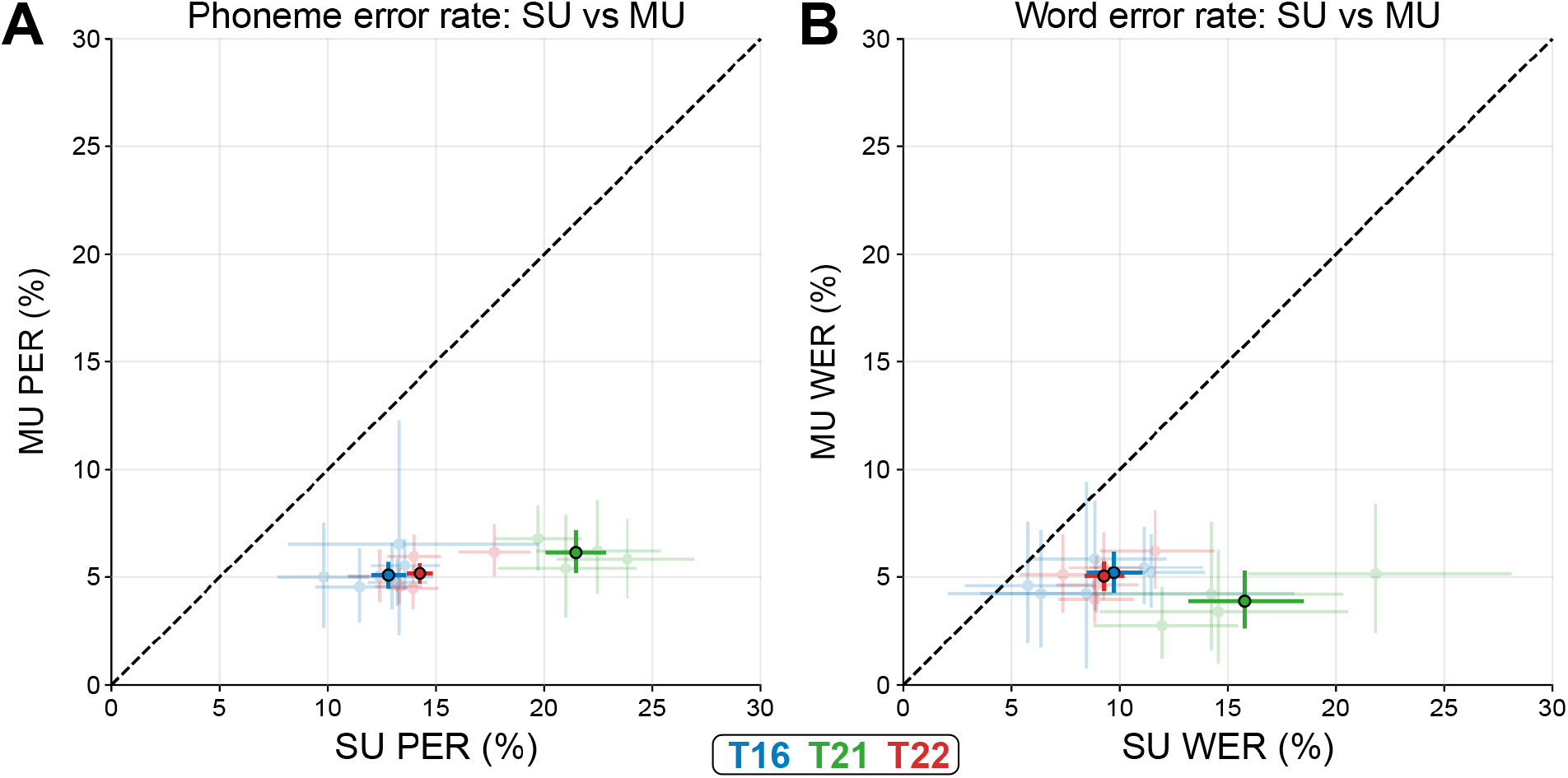
Closed-loop comparison of decoding accuracy between SU and MU models. Phoneme **(A)** and word **(B)** error rates are shown using SU and MU decoding models for closed-loop decoding sessions with participants T16, T21, and T22. Points are aggregate error rates and horizontal and vertical lines are 95% confidence intervals, both colored by participant. Semi-transparent points and lines are individual research sessions, and opaque bolded points and lines are aggregate values across all sessions per participant. Additional details about each session are available in Table S1.

MU models achieved consistently lower phoneme error rates (PER) and WERs than SU models (Figure 2; Table S1). MU models achieved an average 55.9% relative improvement in closed-loop WER over SU models. The accuracy advantage of the MU model also extended to T21’s self-generated sentences, where the MU model achieved an 11.6% WER (95% CI 6.5 - 17.9) compared to a 27.7% WER (95% CI 18.6 - 38.6) with a SU model (Table S3).

### 2.3 Multi-user models improve offline decoding with additional participants

To evaluate the MU model with larger test sets and with data from additional participants (T12, T15, and T17), we performed offline comparisons. We trained SU models for each participant and MU models on data from all six participants using the datasets described in Table 2. Notably, T15’s dataset is very large (∼278,000 sentences), so to also assess the benefit of a MU model for T15 in a data regime that is comparable to that of other participants, we evaluated performance at two dataset sizes: “T15-Small” and “T15-Large”. T15-Large constitutes T15’s entire multi-year dataset^13^, while T15-Small is a subset of ∼11,000 Copy Task trials, identical to the publicly available T15 data released as part of the Brain-To-Text ‘25 Competition^46^. Here, T15-Small serves to exemplify how a multi-user model helps in a moderate-data, high signal-to-noise ratio (SNR) regime. Note that these offline comparisons are not directly comparable to the above closed-loop results due to differences in dataset train/test splits and model training protocols (Section 4.6).

**Table 2.**
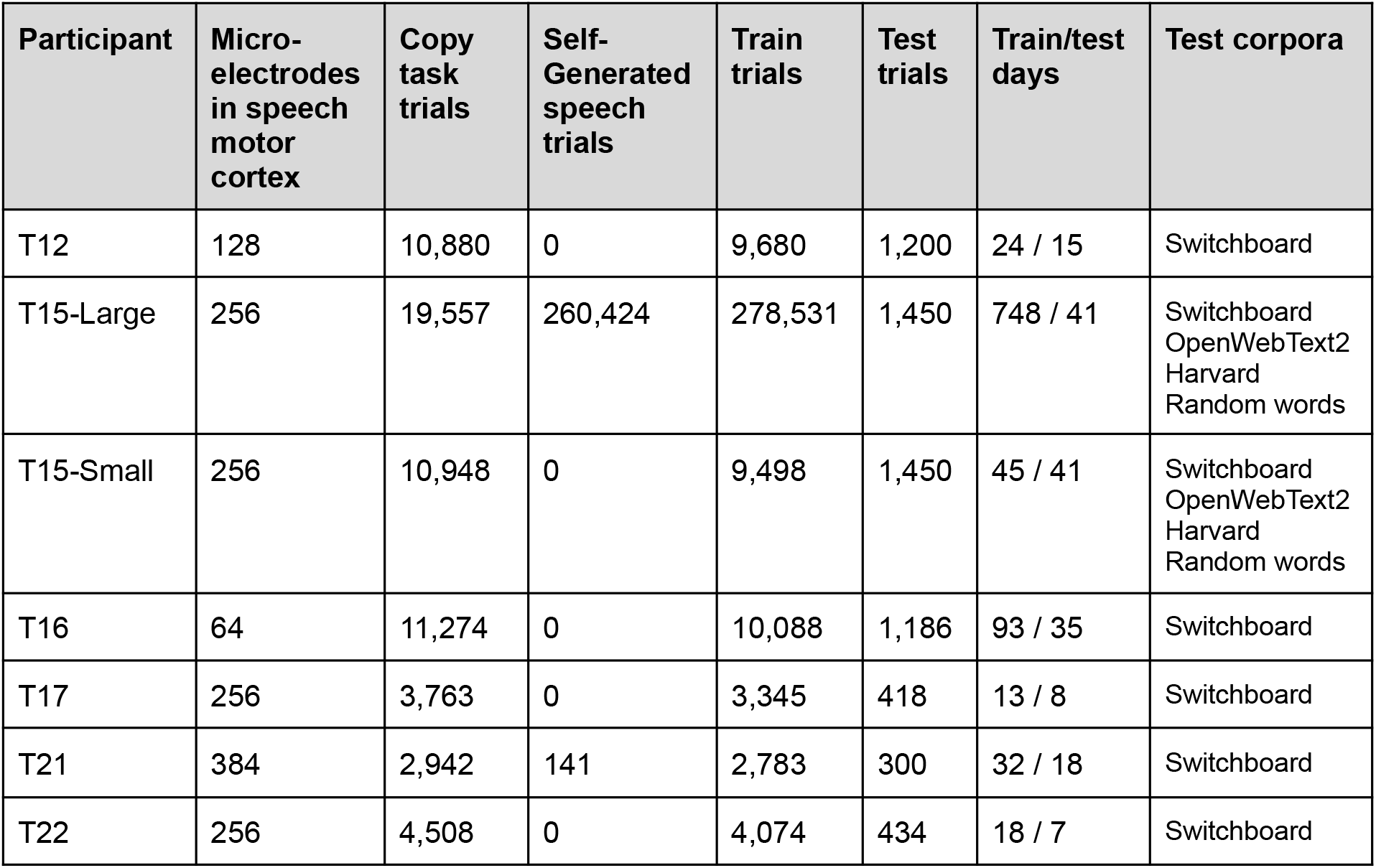
Participant Dataset Information.

MU models outperformed SU baselines for all participants (Fig. 3; Fig. S3). Across the six participants (including T15-Small and excluding T15-Large), the relative WER improvement ranged from 30.1% to 73.3% (mean 51.1%, median 54.2%). T21’s decoding accuracy benefited the most from MU training, with a 73.3% relative WER improvement over the single-user baseline. For T17, who is locked in with anarthria, prior work found large-vocabulary phoneme-based speech decoding mostly unusable as a method of communication (WERs of 39-89%)^19^, consistent with our T17 SU result of 65.6% WER (95% CI 63.9-67.4). However, even in this very low SNR regime, the MU model rescued performance to 23.1% WER (95% CI 21.5 - 24.7), which is comparable to the field’s state-of-the-art as of 2023: 23.8% WER^14^ and 25.5% WER^16^. For T15-Small, the MU model substantially outperformed the SU baseline (30.2% relative WER improvement), demonstrating that MU training is beneficial even when SU models already achieve good decoding accuracy. However, the MU vs. SU performance gap shrank considerably for T15-Large (only 14.3% relative WER improvement), suggesting that MU models may be less advantageous for users with very large training datasets.

**Figure 3.**
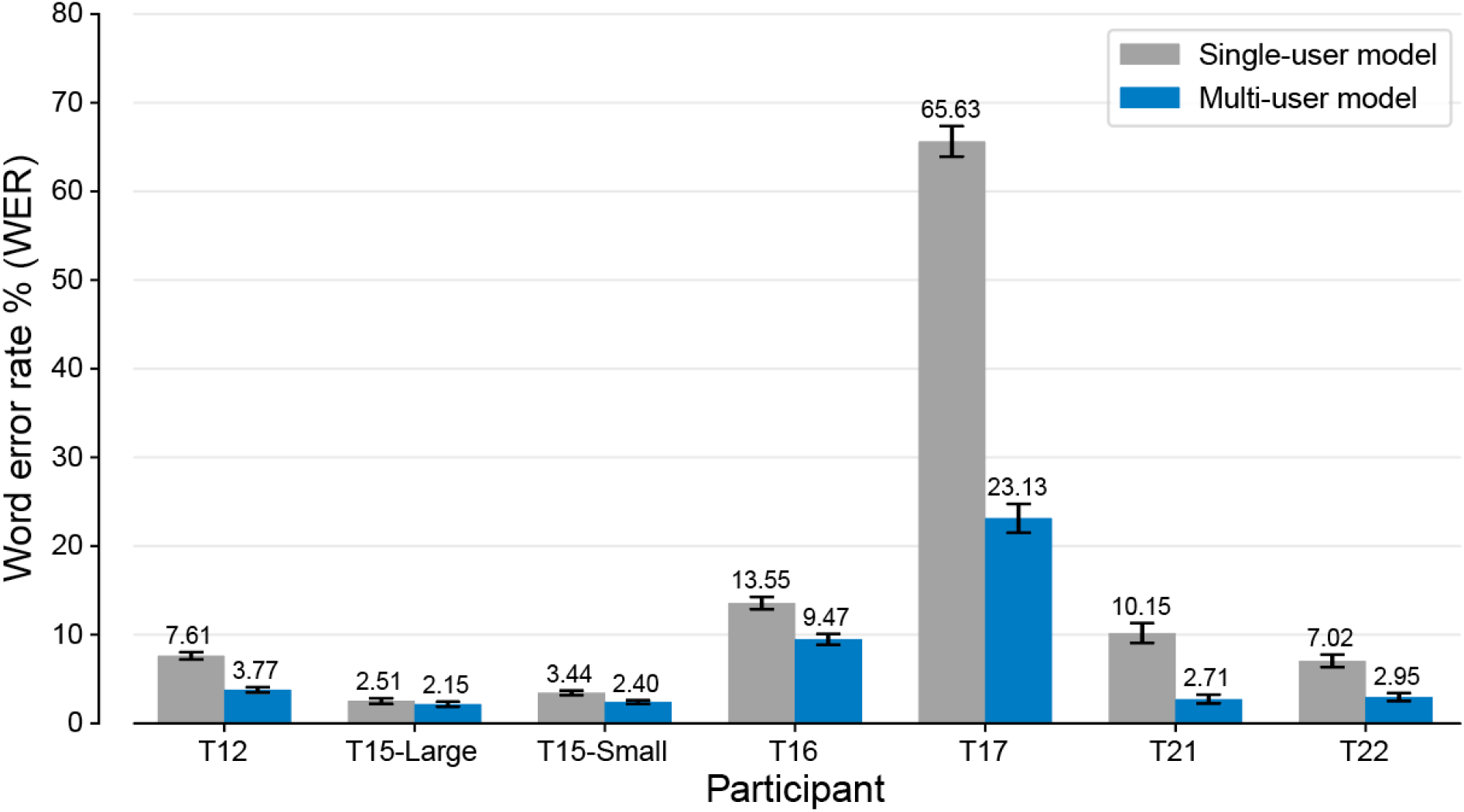
Offline decoding WER comparison between MU and SU models on held-out sentences. MU models were trained on data from all six participants, while SU models were trained only on data from the corresponding participant. For each participant, both models were evaluated on the same held-out test sets (Table 2). The T15-Small MU result is from a model trained using the T15-Small dataset, while all other MU results are from a model trained with the T15-Large dataset. Mean error rates and 95% confidence intervals are an average of three seeds. Phoneme error rates are available in Fig. S3.

### 2.4 Generalizability of the multi-user model scales with more participants

Next, we measured how a MU model’s generalizability to new users scaled with the number of users included in pretraining. We pretrained MU models on pools of one to four participants (amongst T12, T15, T16, T21, and T22) and then evaluated each model on the remaining participant(s) that were excluded from its pretraining pool via frozen-decoder adaptation (Section 4.4.2; Fig. 4A). For frozen-decoder adaptation, a user-projection network was trained to map a held-out participant’s data onto the frozen pretrained MU decoder.

**Figure 4.**
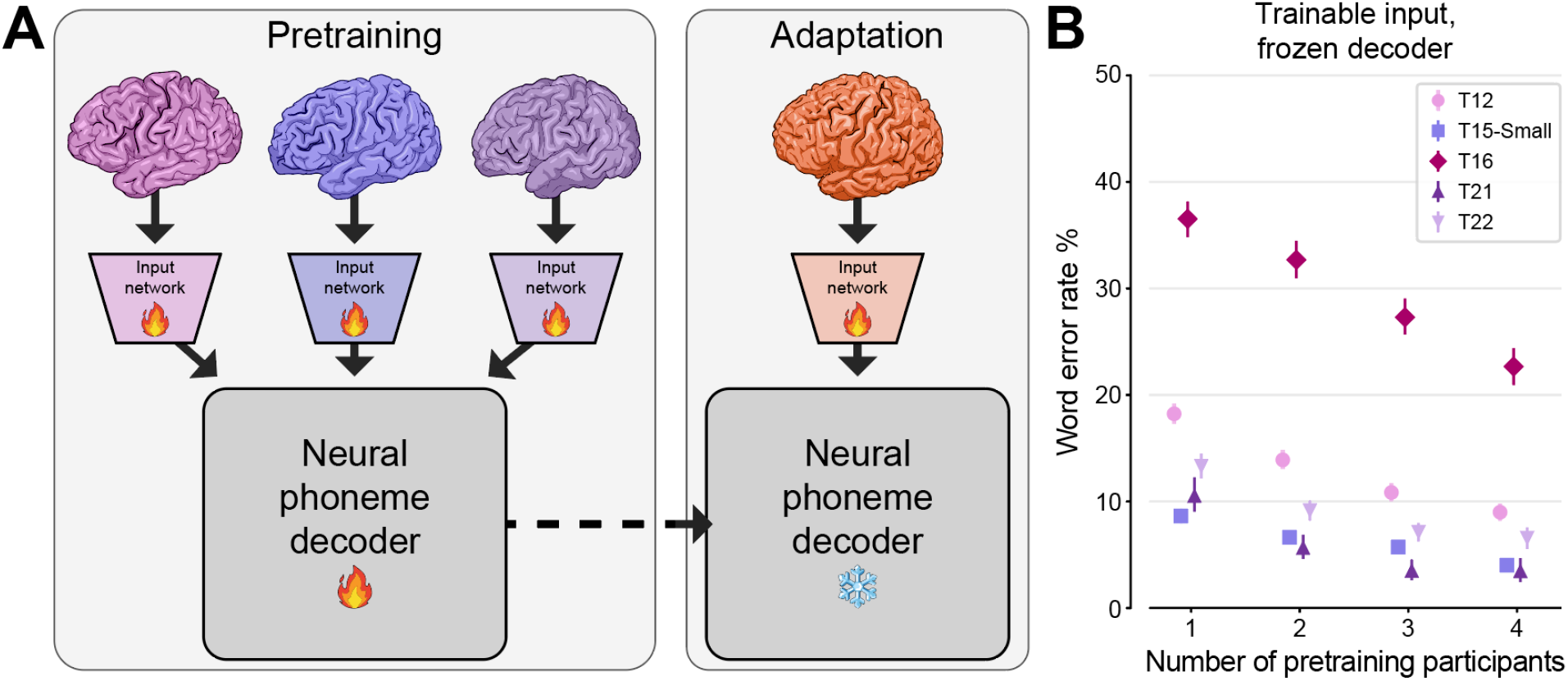
Multi-user model generalization scales with the number of pretraining participants. **(A)** MU models were pretrained on data from 1-4 users, and then adapted to a held-out user (Section 4.6.3). Fire icons indicate trainable network elements, while snowflakes indicate frozen (non-trainable) elements. **(B)** Decoding accuracy for held-out participants after adaptation, plotted as a function of the number of users included in model pretraining. Each point shows the speech decoding accuracy (word error rate) for one held-out participant, averaged over up to 4 models pretrained on different subsets of users, and colored by test participant identity. Error bars denote 95% confidence intervals.

Across all held-out participants, including more pretraining participants improved decoding performance (Fig. 4B). Mapping a held out participant’s data onto the frozen decoder was effective even with a model pretrained on only one participant, and performance improved steadily as the pretraining pool grew. Mean WER (averaged across held-out participants) fell from 17.6% (1 participant) to 9.5% (4 participants). Notably, performance had not saturated at four pretraining participants (17.9% relative improvement in mean WER from n=3 to n=4), suggesting that larger pretraining pools would yield further gains. In the present analysis, we use a relatively expressive user-projection network (a two-layer, feed-forward network), though consistent results were also observed with a simpler user-projection network consisting of a linear projection and tanh nonlinearity (Fig. S6). These results demonstrate that MU models generalize to unseen users, and that this generalization improves as more participants are added to the pretraining pool.

### 2.5 Multi-user models generalize to new users with minimal calibration data

Lastly, we asked whether a MU model could enable new speech BCI users to begin communicating sooner than could be achieved with a SU model. To simulate the decoding performance that a new user would experience on their first day with a speech BCI, we designed an evaluation with the two new BrainGate2 participants introduced in this study, T21 and T22, in which models were trained on randomly selected trials from a single research session and evaluated on a held-out block of trials from the end of that session.

We pretrained two MU models, excluding either T21 or T22, and for each compared two ways of building a usable decoder from the same limited set of trials: few-shot adaptation of the pretrained MU decoder versus training a SU model from scratch. We varied the number of training trials from 10 up to the most each session could support after reserving its final block for testing (up to 90 for T21 and 200 for T22), and repeated this across five research sessions per participant. For few-shot adaptation, the pretrained MU decoder was kept entirely frozen and only a new user-projection layer (learned linear map followed by tanh nonlinearity) was trained (same method as Fig. 4A), restricting adaptation to a minimal mapping onto the model’s existing latent space.

For both T21 and T22, MU model calibration was much more effective than SU training alone. After only 90 training trials — roughly 23 minutes of attempted speech — the MU model reached a WER of 11.2% (95% CI 8.4–14.5) for T21 (Fig. 5A). For T22, the finetuned MU model reached 10.2% WER (95% CI 8.2–12.3) after 90 trials (∼17 minutes of attempted speech) and 6.5% WER (95% CI 4.7–8.2) after 200 trials (∼37 minutes; Fig. 5B). For both participants, most of the MU model’s improvement was realized within the first ∼60-70 trials, with WER continuing to decline more gradually thereafter, indicating that only a small number of trials are needed to map a new user’s neural activity onto the generalizable latent space already learned by the frozen decoder. SU models trained from scratch on the same trials were far less accurate and would not support accurate first-day decoding for either participant: they reached only 87.0% WER (95% CI 85.2–89.2) for T21 after 90 trials, and 76.9% (95% CI 74.8–79.0) and 40.3% (95% CI 37.2–43.3) for T22 after 90 and 200 trials — all far too inaccurate for reliable communication.

**Figure 5.**
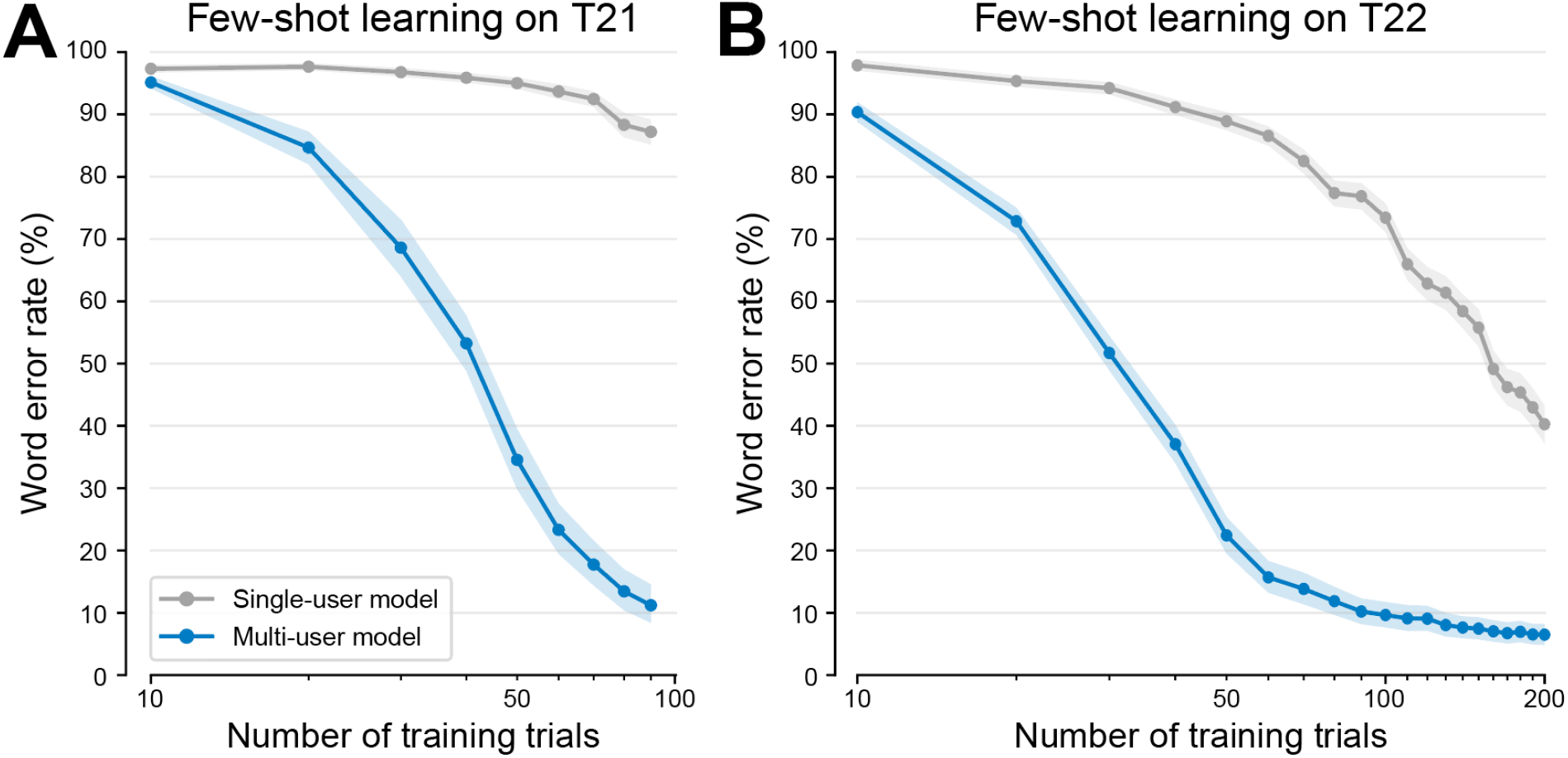
MU models can be rapidly adapted to held-out users via few-shot learning. WER as a function of the number of training trials from limited single-session data using few-shot learning with a pretrained MU model (decoder frozen, only a new participant-projection layer trained) versus a SU model trained from scratch. In both conditions, models were trained on randomly selected trials from a single research session and evaluated on a held-out block of trials reserved from the end of that session. **(A)** Participant T21, decoded with a MU model pretrained on T12, T15, T16, and T22. **(B)** Same as (A) for participant T22, decoded with an MU model pretrained on T12, T15, T16, and T21. Results are averaged across five research sessions per participant, with five random seeds per session. Error bars represent 95% confidence intervals.

These results demonstrate that multi-user pretraining could enable accurate, large-vocabulary speech decoding on a new user’s first day from only minutes of calibration data.

### 3. Discussion

In this work, we introduce a generalizable multi-user speech neuroprosthesis trained jointly across six participants in the BrainGate2 clinical trial. By learning shared phonemic representations from intracortical activity across people who differ in sex, disease etiology, recording array count and locations, and attempted speaking strategy, our multi-user model decoded speech more accurately than single-user models for every participant, both offline and in closed-loop use. Furthermore, multi-user models could be adapted to a new participant with only minutes of calibration data. These results advance the field along three fronts: (1) they replicate and extend high-accuracy intracortical speech decoding across a more diverse participant population, (2) they establish that cross-user generalization substantially benefits intracortical speech BCIs — and in some cases can rescue unusable decoders created from a single participant, and (3) they show that this benefit scales with the number of participants in a way that points toward future decoders that require minimal individual calibration as more participants are added.

### 3.1 Replication and extension across participants

Prior work demonstrating highly accurate, closed-loop, large-vocabulary brain-to-text decoding was largely limited to a single participant (T15)^12,13^, leaving open whether such performance was idiosyncratic to one person’s physiology, array placement, or signal quality. Here we show that comparable accuracy is achievable across participants who span ALS and brainstem stroke etiologies, dysarthria and anarthria, and both vocalized and silent attempted-speech strategies. A MU model trained on left-hemisphere brain signals could even be adapted to right-hemisphere signals of a held-out user (Table S11). That a single jointly-trained model achieves high accuracy across these factors is evidence that accurate intracortical speech decoding is robust and reproducible rather than restricted to one exceptional participant.

### 3.2 Cross-user generalization for intracortical speech BCIs

The success of automatic speech recognition and sEMG interfaces — where training across thousands of users yields models that generalize with minimal calibration^27–30^ — suggested that pooling across users improves decoding performance. However, it is not obvious that success in those domains would transfer to intracortical recordings. Whereas bulk field potential measurements such as EEG, ECoG, or muscle-specific EMG sensor placements might be expected to exploit anatomically-conserved functional tuning across users, the properties of individual neurons in the precentral gyrus are much more heterogeneous^36^ and only loosely follow a somatotopy^34^.

Recent intracortical work found only modest cross-user transfer, confined to low-data regimes^41^. A foundation-model study reported effective cross-user transfer, pretraining a neural encoder across users with a self-supervised masked-reconstruction objective and then finetuning it for phoneme decoding for each individual user^42^. Consistent with this self-supervised objective not yielding task-specific representations, however, their approach still required thousands of labeled trials per user to achieve accurate speech decoding. We find, by contrast, that supervised multi-user training directly on phoneme decoding is itself highly effective — improving accuracy for every training participant and even rescuing usable performance for participant T17, for whom single-user large-vocabulary decoding was previously unusable^19^.

Because this objective drives the pretrained weights to encode robust, user-invariant phoneme representations, adapting to a new participant requires only aligning their neural activity onto the already-learned latent space, enabling few-shot calibration. We attribute our results to two factors. The first is an architecture that pairs a transformer decoder of sufficient capacity to learn shared cross-user representations with per-user input projections; these projections absorb individual differences in neural dimensionality and array placement, forcing the decoder to learn user-invariant phoneme structure. The second is training at a larger data scale than previously used for intracortical speech decoding.

### 3.3 Generalization scales with users and transfers from a shared latent space

The benefit of pooling did not saturate within our cohort. Rather, generalization to held-out participants improved monotonically as the pretraining pool grew, suggesting that including still more participants would yield further gains. This is encouraging given the recent enthusiasm for building large-scale brain foundation models and the anticipated increase in access to human brain data through expanding BCI clinical trials and other intracranial brain recording opportunities.

Furthermore, our adaptation results indicate that this transfer reflects a genuinely shared neural representation. Adapting to a new participant required training only a thin user-projection layer onto a frozen decoder, indicating that the useful phonemic representation already exists in the pretrained weights and a new user need only be aligned onto it. These results suggest that the decoder learns a participant-invariant mapping from neural activity to phonemes — consistent with the neural encoding of attempted speech in ventral precentral gyrus being sufficiently conserved across individuals to support a common decoding model^31–33,41,42^.

### 3.4 Clinical implications

The practical consequence of our work is a substantial reduction in the calibration burden for new BCI users. Previous high-accuracy BCIs required thousands of user-specific sentences before reaching usable accuracy^5,13,14,46^, a time-intensive process that is especially burdensome for people with severe paralysis. For context, modern neuromodulation devices used in everyday clinical practice, such as responsive neural stimulators and deep brain stimulators, require weeks or months of calibration data before providing therapeutic benefit for individual patients^47–49^. By contrast, multi-user pretraining enabled accurate large-vocabulary speech decoding for held-out participants after only minutes of calibration data. This matters most for the participants who can least afford a long calibration: T21, for example, is limited by her ALS to roughly 100 attempted-speech trials per research session — far too few to train an accurate SU decoder, yet sufficient to adapt the multi-user model for usable decoding accuracy. By shortening the path from implantation to reliable communication from weeks of data collection to a single session, multi-user training removes one of the principal obstacles to deploying speech BCIs as assistive devices. We speculate that the same multi-user decoding approach may also serve to rescue decoding performance over longer timescales, where neural signals may degrade due to hardware degradation, immune responses, and/or disease progression, as in ALS^35^.

### 3.5 Limitations and scope

Several limitations qualify our claims. The advantage of multi-user training was smallest for our highest-SNR participant with by far the largest individual dataset (T15), indicating that pooling is most valuable when an individual’s own data is limited or noisy. Our test sentences are drawn predominantly from the Switchboard corpus and reflect conversational English; although we controlled for cross-participant sentence overlap and the multi-user advantage persisted (Table S6), testing with more linguistically diverse material would better establish generality. All participants were implanted with the same microelectrode technology (NeuroPort Arrays; Blackrock Neurotech) in the precentral gyrus; whether the shared representation transfers across recording modalities, array geometries, or implant locations remains to be tested.

### 3.6 Future directions

The lack of MU model saturation at four pretraining users motivates assembling larger multi-user cohorts, with the eventual goal of a model that decodes a new user’s speech with little or no individual calibration, as surface-EMG interfaces now achieve^29^. The shared latent representation may also serve as a foundation for capabilities beyond phoneme decoding, such as multi-user brain-to-voice synthesis^18^ or representations broadly useful for downstream language applications^50^. Architecturally, the cascaded phoneme-then-language-model design we use is effective but may ultimately be surpassed by end-to-end approaches that map neural activity to text directly^42^. More broadly, the demonstration that intracortical speech representations are shared enough to pool across people invites a shift in how the field uses data: rather than treating each participant’s dataset as isolated, pooled multi-user data can be treated as a shared resource that improves every new user’s decoder — a foundation-model paradigm^42,51,52^ for clinical neuroprostheses. In this way, the clinical trial pioneers enrolled in the BrainGate2 clinical trial may now tangibly help future speech BCI users achieve independent communication faster and more effectively than before.

## Acknowledgements

We thank participants T12, T15, T16, T17, T21, T22, and their respective care partners for their immense contributions to this research.

## 4. Methods

All participants in this study were enrolled in the BrainGate2 clinical trial (ClinicalTrials.gov identifier NCT00912041). This pilot clinical trial was approved under an Investigational Device Exemption (IDE) by the US Food and Drug Administration (IDE no. G090003). Permission was also granted by the institutional review boards at the University of California, Davis (protocol no. 1843264), Emory University (protocol no. 00003070), Stanford University (protocol no. 20804), and Mass General Brigham (protocol no. 2009P000505). All participants gave informed consent prior to any experimental procedures. All research was performed in accordance with relevant guidelines and regulations. This manuscript does not report primary clinical-trial outcomes; instead, it describes scientific discoveries that were made using the data collected in the context of the ongoing clinical trial.

This study included six participants (trial designators T12, T15, T16, T17, T21, and T22) with four to six 64-microelectrode arrays (NeuroPort Array; Blackrock Neurotech, Salt Lake City, Utah, USA) implanted in arm-, hand-, speech-, and/or language-related cortical regions (Fig. S1). Four participants have been described in previous studies, while two are introduced here for the first time. For each participant, microelectrode array implant targeting was based on pre-surgical fMRI scan sequences (Human Connectome Project pipelines^53^) that provided estimates of cortical area borders, as in previous speech BCI studies^12,14,19,21^.

### 4.1 Participants

**T12**, a left-handed woman with dysarthria due to bulbar-onset ALS, was 67 years old at the time of enrollment. T12 had two arrays implanted in her left ventral precentral gyrus (area 6v; oral-facial motor cortex) and two placed in her left inferior frontal gyrus (area 44). In this study, we only use neural data from the two area 6v arrays, since the area 44 arrays were previously reported not to have significant contributions to phoneme decoding^14^. See ref. 14 for additional T12 details.

**T15**, a left-handed man with dysarthria due to ALS, was 45 years old at the time of enrollment. T15 had four arrays implanted in his left precentral gyrus; two in ventral 6v, one in area 4, and one in area 55b. In this study, we use neural data from all four arrays. See ref. 12 for additional T15 details.

**T16**, a right-handed woman with dysarthria due to pontine stroke that occurred 19 years prior to enrollment in the clinical trial, was 51 years old at the time of enrollment. T16 had four arrays implanted in her left precentral gyrus; one in area 6v, one on the border of the premotor eye field and area 55b, and two in area 6d. In this study, we only use neural data from the array in area 6v based on prior work showing that the other arrays did not significantly contribute to phoneme decoding^21^. See ref. 21 for additional T16 details.

**T17**, a right-handed man with anarthria due to ALS, was 34 years old at the time of enrollment. T17 had six arrays implanted in his left precentral gyrus; two arrays in area 6d, two arrays in area 6v, and two arrays in area 55b. In this study, we only use the neural data from the arrays in the speech-related areas (6v and 55b). See ref. 19 for additional participant details.

**T21**, a right-handed woman with dysarthria due to ALS, was 57 years old at the time of enrollment. T21 was diagnosed with ALS 3 years prior to enrollment in the clinical trial. Four arrays were implanted in her left precentral gyrus: two arrays in area 6v, and two in area 55b. T21 also had two arrays implanted in her right precentral gyrus, one in area 6v and one in area 55b. In this study, we use data from all six arrays.

**T22**, a right-handed woman with anarthria due to pontine stroke that occurred 3 years prior to enrollment in the clinical trial, was 42 years old at the time of enrollment. She had anarthria and tetraparesis at baseline, and communicated through a combination of silent orofacial gestures and an eye-gaze tracking system. T22 had five arrays implanted in her left precentral gyrus; one in area 6d, three in area 55b, and one in area 6v. T22 also had one array implanted in area 44 of her left inferior frontal gyrus. In this study, we only use neural data recorded from areas 6v and 55b.

### 4.2 Neural recording and feature extraction

Each microelectrode array contains 64 electrodes arranged in an 8×8 grid. Each electrode is 1.5 mm long and measures broadband 30 kHz intracortical activity from a single recording site at the tip. Neural activity was recorded from electrodes using NeuroPlex E headstages (Blackrock Neurotech), connected to percutaneous pedestals that were mounted to the skull. Signals were analog-filtered between 0.3 Hz and 7.5 kHz (4th-order Butterworth), digitized at 30 kHz with 250 nV resolution, and streamed in 1 ms windows to a custom real-time processing node written in Python version 3.8. Each 1 ms window was band-pass filtered between 250 and 5000 Hz using a 4th-order zero-phase Butterworth filter. To reduce edge artifacts, windows were padded using the previous 1 ms of data on the left and mean-padding on the right. Linear regression referencing (LRR)^54^ was applied independently to each array (64 electrodes per group) to suppress common-mode noise.

Two standard neural features were extracted from each electrode: threshold crossing spikes and spike-band power. Threshold crossings were identified when the voltage crossed -3.5 or -4.5 (depending on the participant) × the root mean square (RMS) value for that electrode. Spike-band power was computed by squaring the filtered signal and averaging over the 1 ms window, with a ceiling of 12,500 μV² to reject outliers. This full preprocessing pipeline – including filtering, denoising, and feature extraction – was completed in under 1 ms per window. Extracted features were binned into 20 ms non-overlapping intervals. Binned threshold crossings were computed by summing across consecutive windows; binned spike-band power was calculated by averaging over the same span. Each neural feature was z-scored using statistics from the preceding 5-20 speech trials.

At the beginning of each session, a brief calibration task involving repeated word attempts or silence was used to estimate per-electrode RMS values for thresholding and to compute LRR filter coefficients. These parameters were re-estimated after each block of recording throughout the session to mitigate signal nonstationarities over time.

### 4.3 Speech task design and neural dataset

Neural activity was recorded continuously while the participants attempted to speak sentences in one of two task conditions. Participants used a range of speaking strategies during these tasks, ranging from mimed speech to vocalized speech. We refer to each individual utterance of a performed attempted speaking task as a “trial”. There was no difference in how trials from each task were used for model training.

#### Copy Task

The Copy Task is a structured instructed-delay style research task, where prompted sentences are presented to the participant as text on a screen during a delay epoch. After a “go” cue, the participant is instructed to attempt to speak the presented sentence. For the test datasets in this study, prompted sentences are primarily drawn from the Switchboard corpus^55^. As the participant attempted to speak, they were either shown the decoded words in real-time (“Closed Loop”) or were not shown any feedback (“Open Loop”). Copy Task trials were collected in blocks, each lasting approximately 5 to 30 minutes, where trials were performed sequentially, one after the other; the number of trials performed in one block varied by participant and by research session.

#### Self-Generated Speech

In the Self-Generated Speech Task, the participant said whatever they wanted to and then confirmed the decoding accuracy after each trial, using a custom graphical user interface controlled by an eye-tracker or neural cursor to select an on-screen button. The participants attempted to speak at their own pace, which was typically 30 to 70 words per minute. All self-generated speech trials with T21 were closed loop and collected during research sessions. All self-generated speech trials with T15 were closed loop and collected from both research and independent use sessions^13^. For T21, we included only sentences for which she confirmed the ground-truth label (using the independent use system described in ref. 13). For T15, we additionally included trials that were not initially rated as correct by the participant, but were instead pseudo-labeled using offline retrospective inference with a SU T15 phoneme decoder. These trials were only given pseudo-labels if they could be confidently labeled with a coherent phoneme sequence. Pseudo-labeled trials were only used for training models, and were never used for model evaluation.

Each trial consisted of a time-series of extracted neural features (details in Section 4.2) and a sentence label (the ground truth sentence that the participant attempted to say). Sentence labels for each trial were also converted into phoneme sequences using the g2p-en python package^56^ to provide training labels for the neural phoneme decoder. Any trial longer than 75 seconds was excluded from our dataset to limit GPU VRAM requirements.

We split the neural dataset into training and testing sets for each participant. For T12, we used the training, validation, and testing sets provided in the BrainToText ‘24 Benchmark^57^, combining the original training and validation sets into one larger training set. For T15-Large, we used the testing set provided in BrainToText ‘25 Competition^46^ and trained on all remaining Copy Task and Self-Generated Speech trials. For T15-Small, we used the training, validation, and testing sets provided in the BrainToText ‘25 Competition^46^, combining the original training and validation sets into one larger training set. For T16, T17, T21, and T22, we tested on the last Copy Task block(s) of some research sessions, holding out ∼10% of all sentences for testing.

Since all participants’ training and testing sets contain sentences drawn from the Switchboard corpus, there is some degree of cross-participant sentence overlap. To account for this, we created additional deduplicated training and testing datasets where sentence overlap was controlled for (Table S5) and demonstrated that MU models still outperform SU models for all participants (including T15-Small, but not T15-Large) when trained and tested on these datasets (Table S6).

### 4.4 Multi-user speech BCI

Our brain-to-text speech BCI built upon the real-time architectures described in refs. 12-14, the latter representing the current state-of-the-art. The system was a cascaded multi-stage design and consisted of a neural feature extraction pipeline, followed by a neural phoneme decoder that predicted the most likely phoneme sequences that the user was attempting to say from neural activity, which were subsequently converted into words and sentences by a series of language models (Fig. 1). Words were displayed as text on a screen in real time and could be read aloud by a text-to-speech algorithm at the end of each sentence. For this study, we made advancements to both the neural phoneme decoder and downstream language model pipeline.

#### 4.4.1 Neural phoneme decoder

##### Transformer-based decoder

Our neural phoneme decoder (Fig. 1) is a transformer-based ^58^ decoder that learns a mapping from neural data to phoneme sequences. For the MU model architecture, each participant’s data is projected through a separate input network to the model’s hidden dimension. For a SU model, this participant projection is not used.

After (optional) participant projections, a patch embedding network downsamples the neural data from 20 ms to 80 ms resolution, following refs. 12-14. It concatenates 4 consecutive time-steps, projects the result to the model’s hidden dimension via a learned linear layer, and strides forward by 4 time-steps, repeating across the input. One phoneme prediction is thus made every 80 ms. Patch embedding reduced compute requirements and inference time required to process the neural data, while also improving accuracy. The number of time-steps to concatenate together (patch size) and the number of time-steps to stride forward (patch stride) are hyperparameters of the model (Table S7).

After downsampling, the neural data is processed by a series of multi-headed self-attention transformer decoder blocks. Each decoder block consists of a multi-headed self-attention sublayer followed by a position-wise feed-forward network (FFN), each wrapped in a residual connection. The FFN is a gated linear unit with a SiLU activation^59^: its input is projected up to twice the FFN hidden dimension and split into a value and a gate, the value is scaled element-wise by the SiLU-activated gate, and the result is projected back to the model’s hidden dimension. These decoder blocks iteratively encode the input neural data into contextualized latent representations from which phoneme probability distributions are predicted. Each phoneme probability distribution is over 41 possible classes, which include 39 English phonemes, a silence token, and a blank token for the CTC algorithm^60^. We use Rotary Positional Embeddings^61^ for positional encodings and RMSNorm^62^ for normalizing activations before self-attention and FFN sublayers. We forgo the day-specific neural data transformations that previous speech BCI studies^12–14^ have used, since we found our model robust to neural nonstationarity, so these transformations were not required (Fig. S4).

We train both unidirectional and bidirectional models, which are configured identically, except for the mask applied during self-attention sublayers. For unidirectional models, we apply a causal mask during self attention so that no timestep may attend to any timestep later in the sequence. For bidirectional models we apply only a padding-mask so that timesteps may attend to any other timestep that is not a padding token.

##### Training objective

We trained our model using the CTC algorithm^60^ to map arbitrary-length sequences of neural data to phoneme sequences without requiring a strict alignment between the two. The CTC algorithm maximizes the probability of the true phoneme sequence, marginalizing over all possible alignments of predicted phonemes to the neural data. We also incorporated an auxiliary loss, Intermediate-CTC^63^, which predicts phoneme probabilities from the hidden states of intermediate transformer blocks. These intermediate predictions were computed with the same final normalization and phoneme-prediction projection used for the model’s main output, i.e., the prediction head was shared across the final and intermediate CTC losses. This auxiliary loss encourages intermediate layers to produce phoneme-discriminative representations, improving final-layer accuracy. Intermediate-CTC losses were computed at a subset of transformer blocks spaced roughly evenly through the network, with the specific blocks depending on the model’s depth (e.g., blocks 2, 4, and 6 for an 8-block SU model, and blocks 8, 12, 16, and 20 for a 24-block MU model). We used the PyTorch^64^ CTCLoss implementation to train our models.

##### Regularization

During training we applied stochastic depth^65^ which randomly drops the residual contribution of entire transformer blocks on a per-sample basis. The drop probability increased linearly with depth: block ℓ ∈{1,…,*L*} was dropped with probability 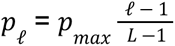, so that the first block was never dropped and the last was dropped with probability *p_max_*. This reflects the intuition that earlier blocks extract low-level features that later blocks depend on, and should therefore be dropped less often. The residual contribution of each surviving block was scaled by 1 / (1 – *p_l_*) so that its expected magnitude matches the test-time value, when all blocks are active. We additionally applied dropout regularization to the outputs of the self-attention and FFN sublayers and to the attention scores.

##### Data augmentation

We applied data augmentation during training to improve the model’s robustness to the variability inherent in neural data and to effectively increase the amount of available training data. Augmentations were applied on-the-fly to each training batch before it was passed to the decoder, and were not applied during validation or inference. We used four augmentations: time-warping, time-masking, and channel-masking (inspired by SpecAugment^66^), and additive white noise (as in previous speech BCI studies^12–14^).

Time-warping locally stretched and compressed each trial along the time axis. Each trial was divided into a small number of contiguous segments by randomly selecting 1 to 5 control points, and each segment was independently stretched or compressed by a factor drawn from a Gaussian centered at 1 (standard deviation 0.1, clamped to the range [0.5, 2.0]). The neural data was then resampled onto the resulting non-uniformly warped time axis by interpolation, which also changed the overall length of the trial.

Time-masking replaced spans of contiguous timesteps, overwriting each timestep with an instance of a learned mask vector. For each batch, we sampled one span length (between 20 and 40 time bins) and one masking probability (between 0.4 and 0.6), both held fixed across that batch’s trials. Then, independently for each trial, we placed masked spans of the sampled length at random start positions, until at least the sampled fraction of the trial’s timesteps had been masked. Because start positions were drawn independently, spans could overlap or extend past the end of the trial, so the realized masked fraction only approximated the sampled probability.

Channel-masking zeroed out both features (threshold crossings and spike-band power) of a random subset of electrodes for the entire duration of a trial. For each batch, we sampled a masking probability (between 0 and 0.1), and then independently zeroed each electrode in each trial with that probability. Finally, we added independent, zero-mean Gaussian white noise (standard deviation 0.5) to every feature at every time step. Exact augmentation hyperparameters are listed in Table S7.

##### Training details

Linear-layer weights were initialized from a normal distribution with standard deviation 1/√(fan-in). Following standard practice for deep residual transformers, the two residual output projections in each block — the attention output projection and the FFN down-projection — were additionally scaled by 1/√(2L), where L is the number of decoder blocks, to prevent the variance of the residual stream from growing with depth^67^.

We trained with mini-batch gradient descent, generally using a batch size of 64. Within each batch, every participant contributes a fixed number of trials (a single participant for single-user models; see Section 4.4.2 and Table S10 for the multi-user case), and a participant’s trials are drawn uniformly at random, with replacement, from their full pool of training trials. All sequences were zero-padded within a batch to the length of the longest sequence. After a batch was sampled, the neural data was augmented (as described above) and temporally smoothed with a Gaussian filter (σ = 40 ms); Gaussian smoothing was applied during both training and inference.

We optimized the model using the AdamW^68^ and Muon^69^ optimizers. All two-dimensional weight matrices were optimized with Muon, with the sole exception of the output phoneme-prediction projection; that projection, together with all remaining one-dimensional parameters, was optimized with AdamW. Weight decay was applied to the weight matrices but withheld from biases, RMSNorm parameters, and the learned mask token. The learning rate followed a cosine decay schedule with linear warmup. Learning rate was scaled down with model depth and volume of training data (Table S7). Before each optimizer step, the global norm of the gradients was clipped to a maximum of 10. Training was performed in automatic mixed precision using bfloat16.

We trained for a fixed number of batches, scaling the number of batches with the size of the training dataset (e.g., 500,000 batches for our 6-user model). Validation on a held-out set and model checkpointing were performed periodically throughout training.

##### Weight Averaging

Instead of selecting the model checkpoint with the best validation set metrics, we employed weight averaging^70^. After a model had been trained (which included periodic model checkpointing after each validation step), we collected the last two thirds of checkpoints, sorted them in increasing order of validation set PER, and then sequentially added the model weights of each checkpoint to a running weight average of the model, keeping a checkpoint only if it did not increase validation PER. The motivation behind this technique is that during the late stages of model training, validation checkpoints tend to oscillate around the same low-error basin in the loss landscape, so a linear interpolation of these checkpoints could lead to lower error than any individual checkpoint. This technique is applied once at the end of model pretraining and results in a single set of weights, so no additional inference or memory costs are incurred during inference of a weight-averaged model.

##### Single-user vs. multi-user models

SU and MU models use the same decoder architecture and training procedure, but differ in how they handle their inputs and in their depth. MU models project each participant’s neural data through a separate, learned linear layer to the model’s hidden dimension; SU models, trained on a single participant, omit this projection. MU models are also deeper: because the MU training dataset is larger and more heterogeneous, they use more decoder blocks (12 - 24, versus 8 for SU models), with the maximum stochastic-depth rate (0.3 vs. 0.2) and the number of Intermediate-CTC layers (four vs. three) increased alongside this added depth. We found that SU models did not benefit from increased model depth or training budget (Fig. S7); hence the depth difference between MU and SU models. Remaining training hyperparameters were scaled with dataset size and are listed in Table S7.

Significant model architecture and training procedure improvements to our neural phoneme decoder that have been introduced since ref. 13 are justified via an ablation study (Table S4). General model architecture and training hyperparameters are included in the Table S7. Our transformer-based neural phoneme decoder scales with dataset size and outperforms the recurrent neural network (RNN)-based decoder used in previous studies^12,14,19,20^ at all dataset sizes (Fig. S5).

#### 4.4.2 Multi-user training

##### Pretraining

When training a MU model, we included trials from all participants in every training batch. This ensures that every gradient update reflects a joint objective over all participants, discouraging updates that reduce loss for one participant at the expense of others. Within each training batch, participants’ trials were sampled roughly in proportion to the size and estimated SNR of each participant’s dataset (Table S10). The exact per-batch ratios were not critical: provided we

trained for enough batches, small differences in each participant’s per-batch trial count had little effect on training dynamics or final model performance, so these ratios were not exhaustively optimized. We did, however, find it generally helpful to sample our highest-SNR participant most heavily (T15, which also contributed by far the most data).

Since the number of recording channels, or “neural dimension”, for each participant varied, the neural data of each participant had to first be projected to a common latent dimension before being collated into a multi-user training batch. For each participant, we learned a projection from their neural dimension (ranging from 128 to 768) to the model’s hidden dimension, which is 512 for all of our analyses. When pretraining a MU model, we found that a linear projection followed by a tanh nonlinearity led to the most generalizable models; we hypothesize that keeping the participant projection minimalist forces the transformer decoder to learn a generalizable phoneme decoding strategy across participants. After neural data passed through its corresponding participant projection network, all samples underwent the same processing steps without any further user-specific logic. When computing mean loss across a training batch, we did not include any participant specific weighting and weighted all samples equally.

##### Adapting to new users

A pretrained MU model can be adapted to a new user — one whose data was not included during pretraining — in one of two ways, differing in whether the pretrained decoder weights are updated. In both cases, a new user-projection network is learned to map the new user’s neural features into the decoder’s shared latent space and every training batch consists of only the new user’s data; user-projection networks for pretraining users are kept frozen. The two approaches differ only in whether the rest of the pretrained decoder is held fixed or allowed to train. The new-user projection network can either be a simple linear projection followed by a tanh nonlinearity (the same type of transformation used for each user during pretraining), or a more expressive two-layer feed-forward network (the same FFN network described in 4.4.1).

###### Frozen-decoder adaptation

The pretrained MU model is kept entirely frozen and only the new user-projection network is trained. This restricts learning to a minimal mapping from the user’s neural space into the shared latent space already learned by the MU model. Because the only trainable parameters are those of the user-projection network, this approach is robust to overfitting and is preferred when calibration data is limited.

###### Trainable-decoder adaptation

The new user-projection network is learned and the pretrained MU decoder is additionally allowed to train, adapting its weights to the new user. To avoid disrupting the pretrained representations and to guard against overfitting on a new user’s data, we unfreeze the decoder gradually rather than all at once: the decoder is first held frozen while only the user-projection network is trained, and after a fixed number of steps the patch embedding network and earliest decoder block are unfrozen, with successive blocks unfrozen from earliest to latest as training progresses. Allowing the decoder to adapt in this way can yield further accuracy gains when sufficient calibration data is available for the new user.

#### 4.4.3 Phoneme to words language model pipeline

After the neural phoneme decoder produces a sequence of per-bin phoneme probabilities, these are passed to a two-stage language model that converts them into word sequences. In the first stage, a 5-gram language model performs a CTC beam search over the phoneme probabilities to produce an n-best list of candidate sentences, scored by a combination of the acoustic (decoder) score and the n-gram language model score. The top candidate is displayed on screen in real time as the participant speaks. In the second stage, at the end of each sentence, the n-best candidates (up to 100) are rescored by a large language model (LLM), and the highest-scoring sentence after rescoring is displayed as the finalized output. The 5-gram model was trained on the OpenWebText2 corpus^71^ with a 125,000-word vocabulary derived from the CMU Pronouncing Dictionary^72^. We used one of two interchangeable implementations of this pipeline, described below, which achieve comparable decoding accuracy (Fig. S2).

#### WFST implementation

The first 5-gram model implementation is the weighted finite state transducer (WFST) decoder used in prior work^12–14,19^, built on Kaldi^73^ and OpenFST. The phoneme-to-word search is compiled into a single TLG (token-lexicon-grammar) finite-state transducer, and beam search is performed over this graph; a separate unpruned grammar FST is used for lattice rescoring. The n-best candidates are then rescored with the Facebook OPT-6.7B LLM^74^, loaded in 16-bit precision (∼13 GB VRAM). The full unpruned 5-gram model requires approximately 350 GB of RAM to load, representing the vast majority of memory usage of the entire BCI. This language model implementation can be found at: https://github.com/Neuroprosthetics-Lab/nejm-brain-to-text

#### Flashlight-Text/KenLM implementation

The second is a new pipeline, introduced here, built on KenLM^44^ and the Flashlight-Text LexiconDecoder^43^. It uses the same underlying 5-gram language model but achieves equivalent accuracy with substantially lower computational requirements (Fig. S2): it loads the n-gram model as a memory-mapped trie rather than holding the full model in RAM (<10 GB RAM at runtime versus ∼350 GB; <10 s versus ∼15 min load time) and decodes >3× faster (Fig. S2), making it suitable for deployment on a single consumer GPU. The beam search scores each hypothesis as a weighted sum of the acoustic score, the n-gram score, and a per-word penalty. The n-gram scores could also be biased toward certain hotwords (e.g., names) to boost the scores of beams that contained those words, enabling user-specific personalization. The n-best candidates are rescored with a more modern, lighter-weight LLM (Qwen3.5-4B^45^, ∼8–10 GB VRAM in bfloat16), with each candidate scored by the sum of its per-token log-probabilities and combined with the beam-search score via a tunable weight. The lower per-trial inference latency of this pipeline additionally allowed us to compute word-level validation WER during decoder training (rather than relying on phoneme error rate as a proxy) and to perform efficient Bayesian (Optuna TPE) hyperparameter optimization over the decoder parameters. The rescoring LLM can also optionally be finetuned with low-rank adaptation (LoRA) on domain-specific text to further improve rescoring. Decoder and beam-search hyperparameters are listed in Table S12. This novel language model implementation can be found at: https://github.com/Neuroprosthetics-Lab/phoneme-to-words-lm

#### 4.4.4 Continuous online finetuning

While all neural decoding models used in this study were initially pretrained offline with one or more participants previously collected data, models that were evaluated online with a participant were continuously finetuned during real-time use. This enabled models to adapt to day-specific neural variability by updating weights based on new trials from the current day, along with older data drawn from a replay buffer, increasing decoding stability over time. Online finetuning was originally introduced for a handwriting BCI^75^, and a similar approach was adapted for speech decoding ^12,13^. Here, we used a modified version of the speech decoder finetuning protocol described in ref. 13. Key modifications include (1) maintaining replay buffers of intermediate days of data between the original pretrain data and the current day’s data, which improved decoding stability over long periods as in Fig. S4, and (2) integrating the KenLM language model pipeline such that WERs could be calculated for validation data, avoiding overfitting to validation PER.

During online inference, finetuning began once at least six trials from the current day had been collected. Each new trial had a 66.7% chance of being added to the training buffer and a 33.3% chance of being held out for validation. Training and validation buffers were maintained throughout the day. Once sufficient data had accumulated, finetuning proceeded for up to 300 - 500 batches using a cosine learning rate scheduler (no warmup). Each batch (batch size = 64) included 16 trials drawn from four days: the current day and three additional randomly selected days. Every 10 batches, the model was evaluated on the entire validation buffer, and a new checkpoint was saved if the aggregate validation PER improved. At the start and end of each trial, the most recent checkpoint was loaded for inference.

#### 4.4.5 Closed-loop inference

During closed-loop speaking epochs, neural signals were decoded into text and displayed on-screen in real time using a causal neural-to-phoneme model. At the end of each sentence, text was re-predicted using a bidirectional neural-to-phoneme decoder model, candidate sentences were rescored with an LLM, and the final decoded sentence was displayed on-screen and read aloud via text-to-speech. This two-stage setup provided the participant with real-time feedback from the causal model during attempted speech while still benefiting from the bidirectional model’s improved accuracy at the end of each sentence, with minimal added latency. Both the causal and bidirectional phoneme decoders were continuously finetuned in the background on data from the current day, mixed with previous days’ data from a replay buffer to prevent catastrophic forgetting, consistent with previous studies (Section 4.4.4).

### 4.5 Offline finetuning simulation

For offline analyses, we simulated online finetuning to replicate real-time usage. Pretrained models were loaded and periodically updated using the same procedure described in Section 4.4.4: every *n* trials, finetuning was triggered, followed by *n* additional trials of inference before repeating (*n* varied by analysis, see supplement for details). This approach enabled us to approximate the online finetuning dynamics and achieve results consistent with those observed in real-time inference (Table S2). The complete offline continuous finetuning codebase used in this study is included with this manuscript. Hyperparameters and details on the computational resources required for offline continuous finetuning are included in the supplement.

### 4.6 Analysis Procedure

All offline analyses were conducted with bidirectional neural phoneme decoders since real-time decoding was not a factor. All offline analyses used the Flashlight-Text/KenLM language model implementation (Section 4.4.3).

#### 4.6.1 Closed-loop SU vs. MU model comparison (Fig. 2)

We compared SU and MU decoding accuracy in closed-loop research sessions with three participants (T16, T21, and T22; Table S1). In each session, the participant attempted to speak prompted Switchboard sentences in a Copy Task while a single model type — SU or MU — decoded in real time. SU models were pretrained on all available data for that participant and then continuously finetuned throughout the session (Section 4.4.4). MU models were pretrained on all available data from the other participants, adapted to the session participant’s preexisting dataset via trainable-decoder adaptation, and likewise continuously finetuned throughout the session.

Because each session used only one model type, we obtained the counterpart model’s accuracy on the same session data using an offline re-inference pipeline that replicated online inference and finetuning (Section 4.5). This pipeline closely reproduced online accuracy for T15, T21, and T22 (Table S2), validating its use for direct SU vs. MU comparison. For T16, however, the offline pipeline consistently outperformed the online results; we traced this to bugs in the real-time inference code used during T16’s sessions. Because the pipeline reliably reproduced online accuracy for the other participants, we used it to compute both the SU and MU accuracies reported for T16. Thus, in Fig. 2, for T21 and T22 we pair each session’s real closed-loop MU accuracy with the SU accuracy from offline re-inference; for T16 we report both model types from offline re-inference (Table S1).

Per-participant aggregate phoneme and word error rates were computed by pooling phonemes and words (and their respective errors) across all of a participant’s sessions. We then averaged the three participants’ aggregate WERs to obtain the reported 55.9% relative MU improvement. T15 was excluded from this closed-loop comparison because his dataset is orders of magnitude larger than the others’; Fig. 3 instead examines the effect of dataset scale directly via the T15-Small and T15-Large comparison.

#### 4.6.2 Offline single- vs. multi-user comparison (Fig. 3)

We compared offline decoding accuracy between SU and MU models on held-out test sentences for all six participants (Fig. 3). For every participant, a SU model trained on only that participant’s data and a MU model trained jointly across participants were evaluated on the same held-out test set (Table 2); we report both PER and WER.

##### Training

All models were trained on the datasets in Table 2 using the architecture and training hyperparameters in Table S7. Each SU model was trained on a single participant’s data; for T15 we trained two SU models, one on the T15-Small dataset and one on the T15-Large dataset, corresponding to the two T15 data regimes. We trained two MU models. The first was trained on T12, T15-Large, T16, T17, T21, and T22 and supplied the MU result for each of these participants (including the T15-Large bar in Fig. 3). The second substituted T15-Small for T15-Large while keeping the other five participants fixed, and was used only for the T15-Small MU result. This dedicated T15-Small MU model ensures the T15-Small MU-vs-SU comparison reflects only data available in the T15-Small regime, rather than benefiting from the far larger T15-Large dataset entering the pretraining pool. MU models drew per-batch participant trials at the sampling rates listed in Table S10.

##### Evaluation

Each trained model was evaluated statically on its participant’s held-out test set (Table 2): the model was applied directly to the test trials with no continuous finetuning or further adaptation.

##### Aggregation and Statistics

We trained three seeds per model. Following Section 4.8, we first averaged each test trial’s edit errors across the three seeds, then aggregated across trials to obtain mean PER and WER, with 95% confidence intervals estimated by bootstrap resampling. Each bar in Fig. 3 is one participant’s seed-averaged error rate for one model type. For each participant we also computed the relative WER improvement of the MU model over the SU model as (WER_SU − WER_MU) / WER_SU. We report the mean, median, and range of these per-participant relative improvements across participants. T15-Large was excluded from this summary so that T15 contributes only once via T15-Small.

#### 4.6.3 Scaling generalization with pretraining participant count (Fig. 4)

To measure how generalization to unseen users scales with the size of the pretraining pool, we pretrained MU models on between one and four participants and evaluated each on the held-out participant(s) absent from its pretraining pool via frozen-decoder adaptation (Section 4.4.2). This analysis used participants T12, T15, T16, T21, and T22. T17 was excluded from this analysis due to his signals’ much lower decoding accuracy compared to all of the other participants, such that we considered the T17 data to be in a very different SNR regime where transfer learning to the other participants was unlikely to be feasible.

##### Pretraining

For each pool size n ∈ {1,2,3,4} we pretrained five MU models, with participant combinations chosen so that each of the five participants appeared in the pretraining pool exactly n times across the five models. Equivalently, each participant was held out from exactly 5 - n of the five models at each pool size (e.g. held out from all four other models at n = 1, and from one model at n = 4). Unlike our other MU models, each training batch sampled all pretraining participants in equal proportion rather than by data size and SNR. In an attempt to isolate pool size as the only variable, decoder depth was fixed at 16 blocks (Intermediate-CTC calculated at blocks 4,8,12) regardless of n, and the training budget was scaled with pool size to control for overfitting on smaller pools; 100k, 200k, 400k, and 500k training batches for n =1,2,3, and 4. A one-user 16-block model trained for 100k batches is architecturally identical to a 16-block SU model trained for the same budget (apart from the user-projection network); Fig. S7 shows that such a model is not meaningfully overfit, confirming that lower performance at small pool sizes reflects limited cross-user transfer rather than overfitting. To limit compute requirements, this analysis used a patch size and stride of 5 and a batch size of 48. MU models whose pool included T15 used the T15-Large dataset.

##### Evaluation

Each pretrained MU model was adapted to each of its held-out participants with frozen-decoder adaptation. The pretrained decoder was frozen and a new user-projection network was trained on the held-out participants’ full training set (Table 2; T15-Small when T15 was held out), and then evaluated on that participants’ held-out test set. We tested using both a linear projection + tanh (Fig. S6) and a two-layer FFN (Fig. 4B). The FFN used the same module as the decoder blocks (Section 4.4.1), with input dimension equal to the held-out user’s neural dimension, intermediate dimension 512, and output dimension 512 (the decoder hidden dimension). Per-participant adaptation training hyperparameters (e.g. training batches, data augmentation parameters, etc.) are the same as those listed for corresponding SU models in Table S7 except that AdamW learning rate max and Muon learning rate max were 2e-4 and 6e-4 respectively.

##### Aggregation and Statistics

At pool size n, a given participant was held out from 5 - n models, yielding 5 -n independent WER evaluations on their single fixed test set. As in Section 4.8, we first averaged each test trial’s edit errors across these evaluations, then aggregated across trials to obtain mean WER and estimated 95% CIs. Each point in Fig. 4 is one held-out participant at one pool size.

#### 4.6.4 Few-shot learning of held-out participants (Fig. 5)

To test whether a pretrained MU model can be rapidly adapted to a new participant — one whose data was excluded from pretraining — from only a single session’s worth of calibration data, we compared two ways of building a usable decoder from a limited number of trials: few-shot adaptation of a pretrained MU model versus a SU model trained from scratch (Fig. 5). We performed this analysis for T21 and T22, evaluating decoding accuracy as a function of the number of calibration trials.

##### Pretraining

For each target participant we pretrained one MU model whose pretraining pool excluded that participant: the MU model for T21 was pretrained on T12, T15, T16, and T22; the MU model for T22 was pretrained on T12, T15, T16, and T21. Both used the T15-Large dataset for T15 and followed the architecture and training hyperparameters in Table S7 except for the number of decoder blocks being 16 (Intermediate-CTC loss calculated at blocks 4,8,12) and patch size and patch stride being equal to 5. Per-batch participant trials were equal proportions.

##### Few-shot adaptation and SU baseline

From each pretrained MU model, we built a decoder for the held-out participant via frozen-decoder adaptation (Section 4.4.2): the pretrained MU decoder was kept entirely frozen and only a new user-projection network — a learned linear map followed by a tanh nonlinearity — was trained, restricting adaptation to a minimal mapping of the new participant’s neural features onto the latent space already learned by the MU decoder. Few-shot adaptation hyperparameters are listed in Table S9. As a baseline, we trained a SU model from scratch on the same trials. Because the single-session training sets used here are far smaller than the full-participant datasets used elsewhere, the SU models used a scaled-down configuration tuned for these smaller datasets (4 decoder blocks; full architecture and training hyperparameters in Table S8); this configuration was held fixed across all training-set sizes. SU models for this analysis also used a patch size and patch stride of 5.

##### Single-session training and test sets

For each participant we analyzed five research sessions, treating each session independently. Within a session, we reserved the final block of trials as a fixed held-out test set — 15 trials for T21 and 40 trials for T22 — and drew all training trials from the trials preceding that block. We varied the number of training trials in increments of 10, from 10 up to the most each session could support after reserving its test block: 10–90 trials for T21 and 10–200 trials for T22. At each training-set size, training trials were drawn at random, without replacement, from this pre-test pool, with an independent draw for each seed; the same reserved test block was used for every training-set size within a session.

##### Quantifying calibration time

We quantified the amount of attempted-speech data in a training set as the total duration of neural data trained on, computed by summing the number of 20 ms bins across all trials in the training set.

##### Aggregation and statistics

We trained five random seeds per training-set size for each session and participant. For each session and training-set size, we first averaged each test trial’s edit errors across the five seeds, then pooled the seed-averaged test trials across all five sessions and aggregated to obtain a mean WER at that training-set size, with 95% confidence intervals estimated by bootstrap resampling (Section 4.8). Each point in Fig. 5 is the resulting aggregate WER for one decoder type (few-shot MU or from-scratch SU) at one training-set size.

### 4.7 Evaluation

Consistent with prior work, we evaluate brain-to-text decoding performance using PER and WER, both computed via Levenshtein distance. This metric reflects the number of insertions, deletions, and substitutions required to transform the decoded sequence into the ground truth, applied at either the phoneme or word level. We measure words per minute (WPM) as the number of words spoken in a trial divided by the minutes elapsed between the first and last phoneme predictions of a trial. Certain trials were excluded from evaluation if the prompted sentence included a typo or if the participant did not attempt to speak.

### 4.8 Statistics

Word and phoneme error rates were aggregated across trials by summing all edit errors and dividing by the total number of target phonemes or words. Confidence intervals were estimated using bootstrap resampling: individual trials were resampled with replacement (10,000 iterations), and aggregate error rates were recalculated for each resample to generate the empirical distribution. In analyses where multiple model seeds were used to calculate error rates and confidence intervals on the same fixed test sets, edit errors were trial-averaged across seeds before aggregation across trials.

### 4.9 Compute resources

#### Offline analyses

Offline analyses and model training were conducted on six machines running Ubuntu 22.04 LTS, each with one or two GPUs (Nvidia RTX 4090s, 5090s, or 6000 Pro Blackwells). Each computer had between 128GB and 512GB of system memory.

#### Online research sessions

The real-time speech BCI was implemented using a local computer setup that contained ample GPU, CPU, and memory resources to enable high-bandwidth neural recording and low-latency neural decoding. This local setup interfaced with the NeuroPlex E headstages to control neural recording, processed raw 30 kHz neural data, and extracted neural features. This setup was also responsible for CPU- and GPU-intensive operations including real-time decoder inference, phoneme-to-word language modeling, online decoder finetuning, and user-facing task display. Each participant’s setup varied physical hardware and divided processes across different machines. All machines were connected via a local area network and synchronized using the Backend for Realtime Asynchronous Neural Decoding (BRAND) framework^76^. Code was implemented in Python, C, and MATLAB.

---

**Figure S1.**
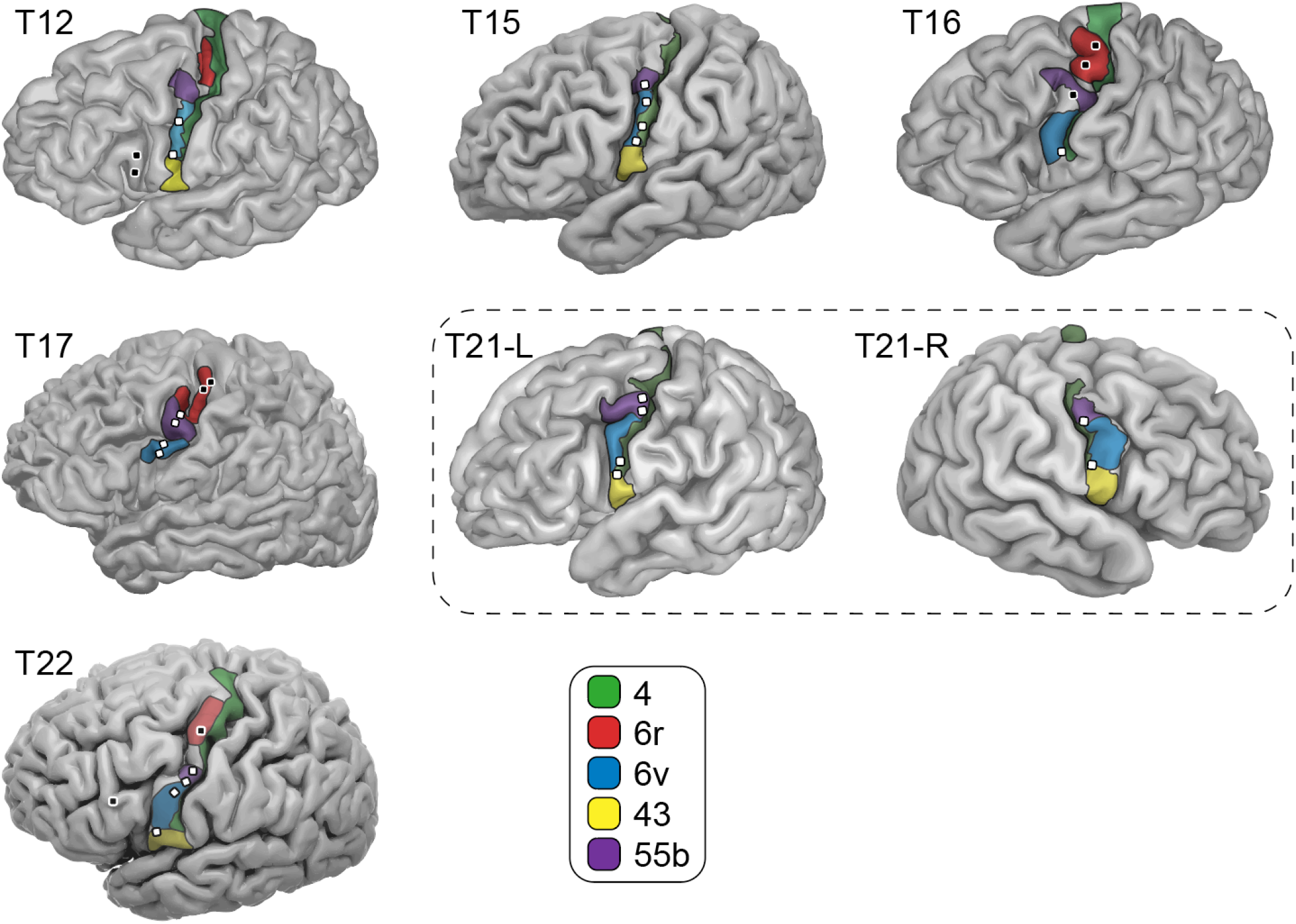
Microelectrode array locations for each participant. Each participant’s cortical structure was reconstructed from pre-surgical fMRI scan sequences. Estimates of relevant cortical areas (acquired through the Human Connectome Project pipelines) are overlaid on each brain. Overlaid white and black squares denote approximate implanted microelectrode array locations. Speech-relevant microelectrode arrays (Table 1) are white with a black outline; data from other microelectrode arrays (black with white outline) were not used in this study.

**Figure S2.**
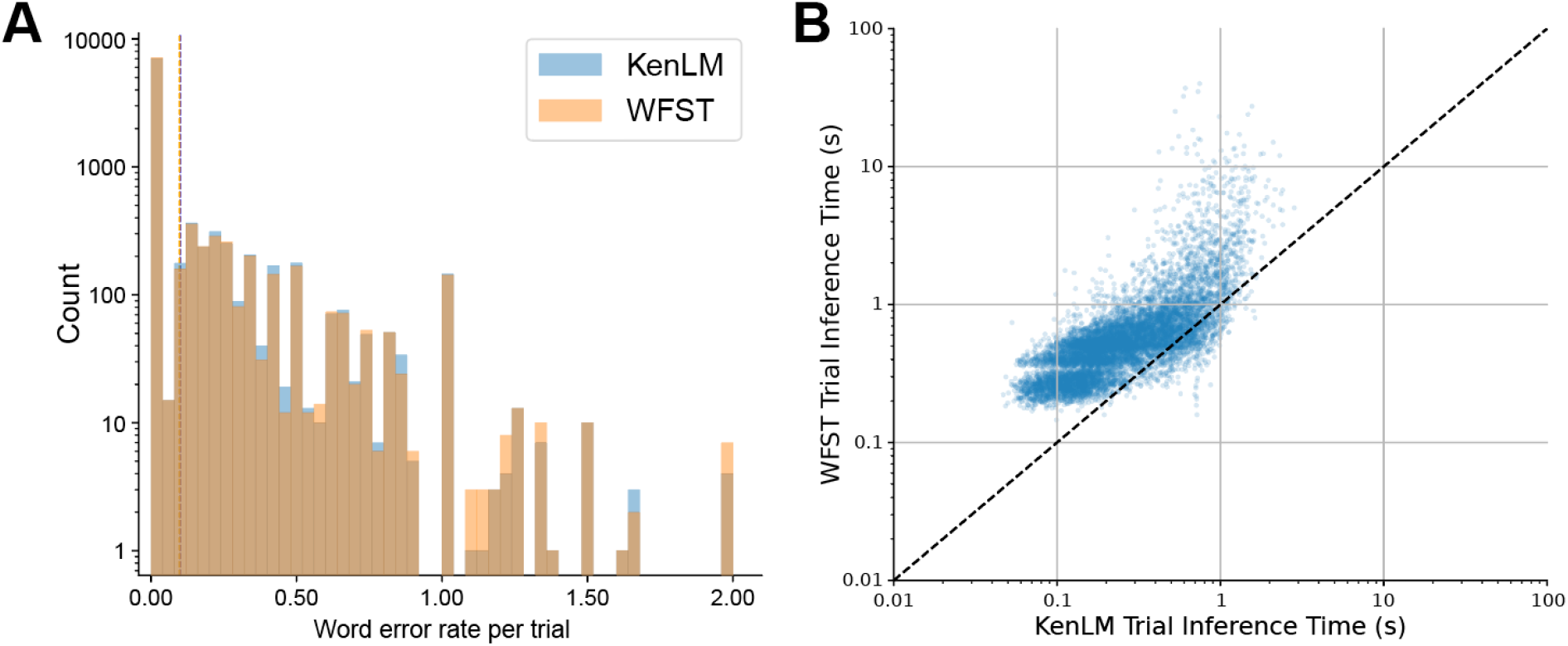
Comparison between WFST and KenLM language model pipelines. Both language model pipelines were evaluated on SU and MU model phoneme logits from all held-out test trials in Fig. 3 (9,616 total trials), which were sourced from all six users. **(A)** Word error rate distribution for both language model pipelines amongst all test sentences. The WFST pipeline produced sentences with an aggregate WER of 8.81% (95% CI: 8.43 - 9.20), and the KenLM pipeline produced sentences with an aggregate WER of 9.08% (95% CI: 8.68 - 9.47). **(B)** Per-trial inference latency for each of the two language model pipelines. The WFST pipeline had a mean inference latency of 0.770 seconds (95% CI: 0.742 - 0.798), and the KenLM pipeline had a mean inference latency of 0.316 seconds (95% CI: 0.310 - 0.321). On a trial-by-trial basis, the KenLM pipeline was an average of 3.11 times faster than the WFST pipeline.

**Figure S3.**
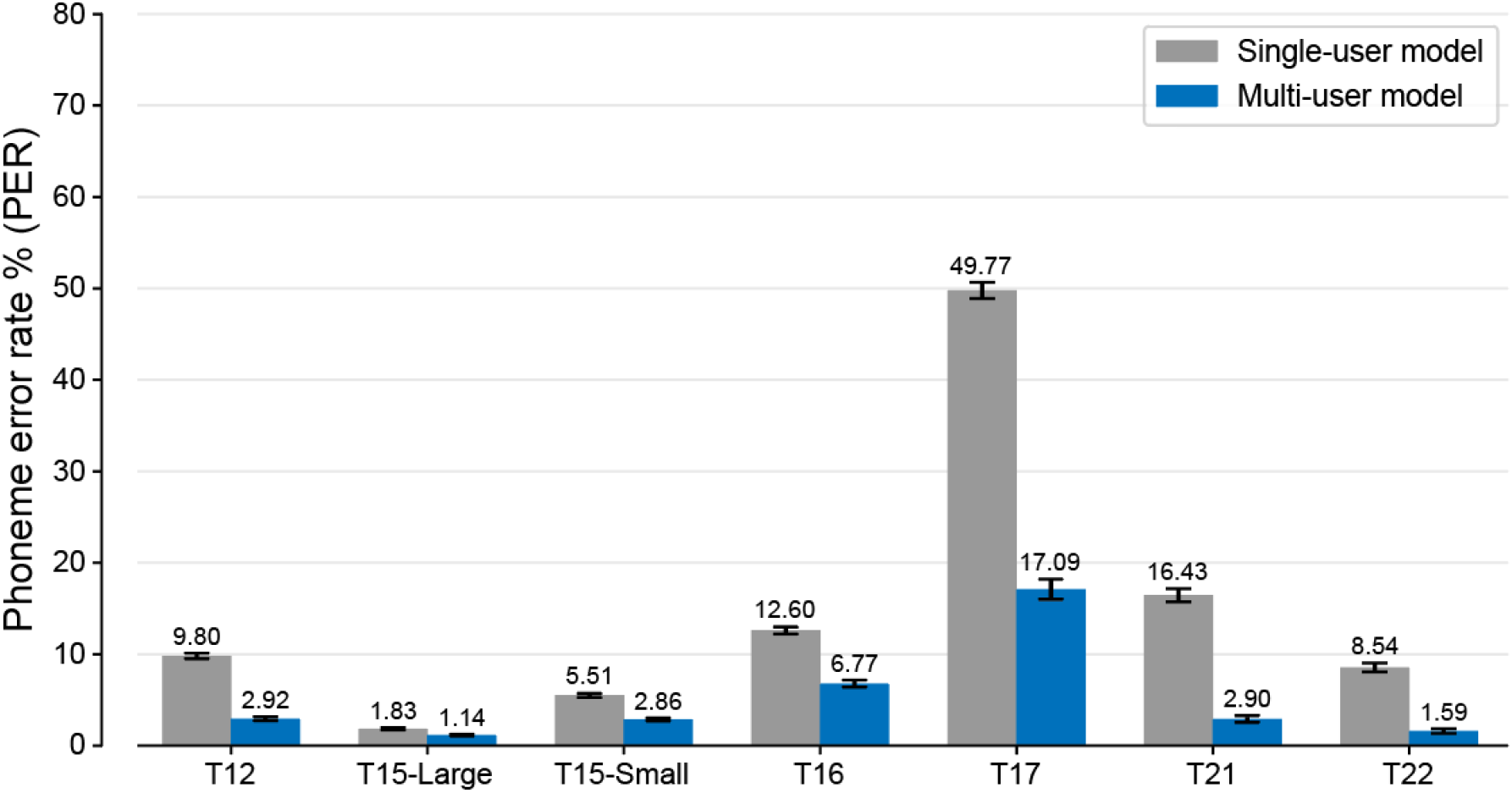
Offline decoding PER comparison between MU and SU models on held-out sentences. The same analysis as in Fig. 3, but here showing phoneme error rates (rather than word error rates as in Fig. 3).

**Figure S4.**
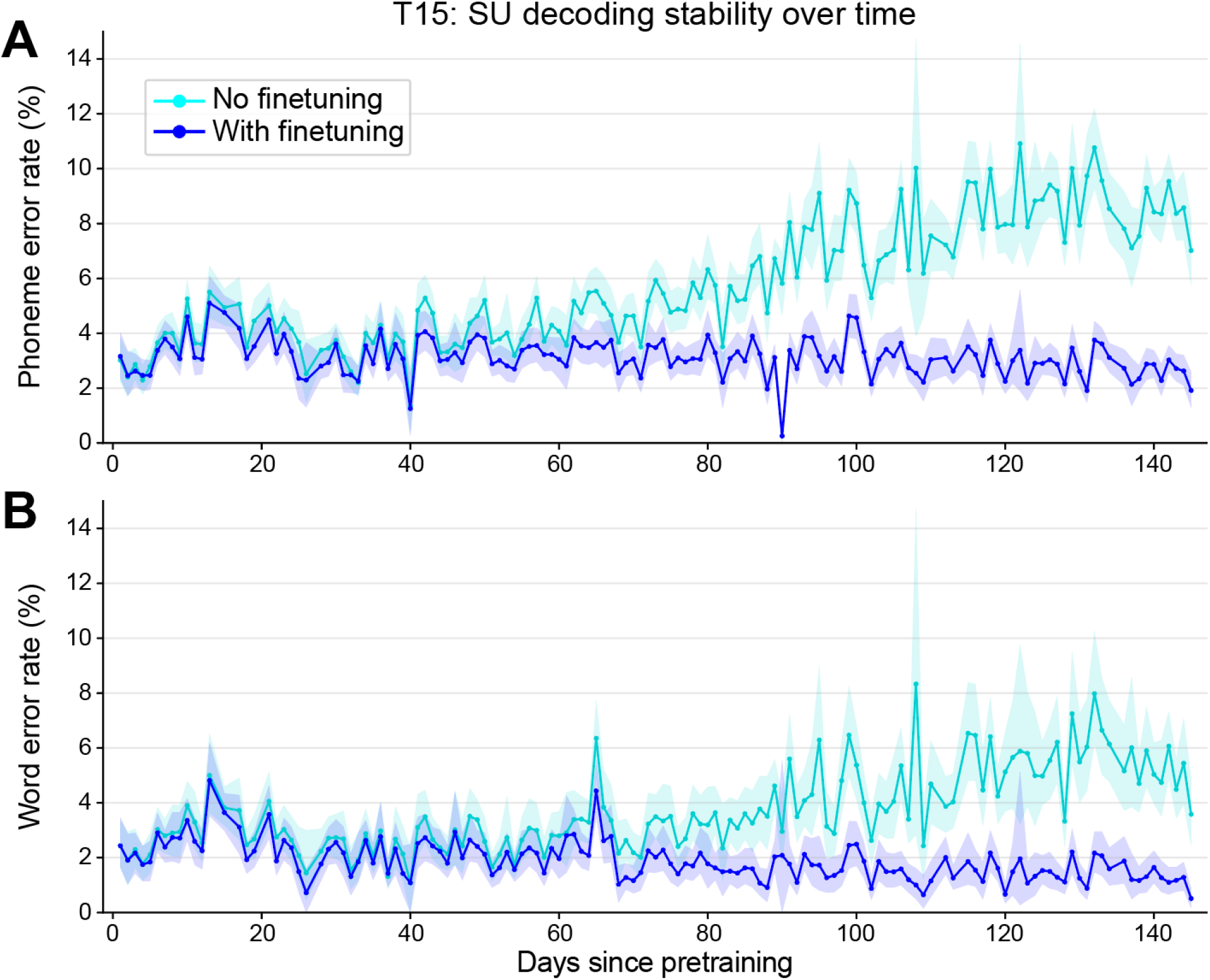
Offline SU model stability dynamics. PER (top) and WER (bottom) as a function of the number of days between model training and evaluation. Models were trained on data from the first 322 days of the T15-Large dataset and evaluated on the next 143 days (including both Copy Task and Self-Generated Speech Task trials) with and without continuous finetuning. Each point represents the aggregate PER across all sentences in a single day. Shaded error bars indicate 95% confidence intervals.

**Figure S5.**
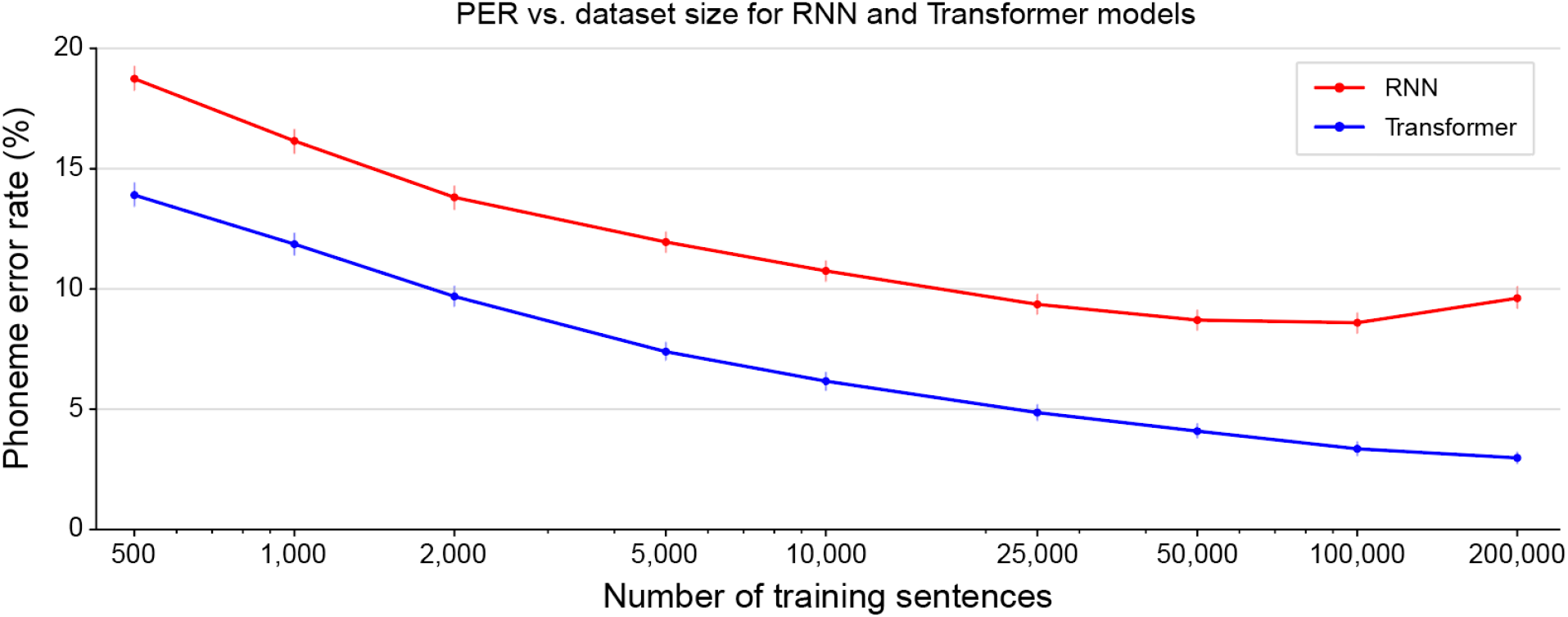
Effect of pretraining dataset size. PER on six held-out days of T15 Copy Task data using continuous finetuning as a function of the number of training sentences (from adding sequential days of data) to pretrain a T15-Large single-user model, either with a recurrent neural network (RNN)-based neural phoneme decoder, or our transformer-based neural phoneme decoder. Error bars are 95% confidence intervals.

**Figure S6.**
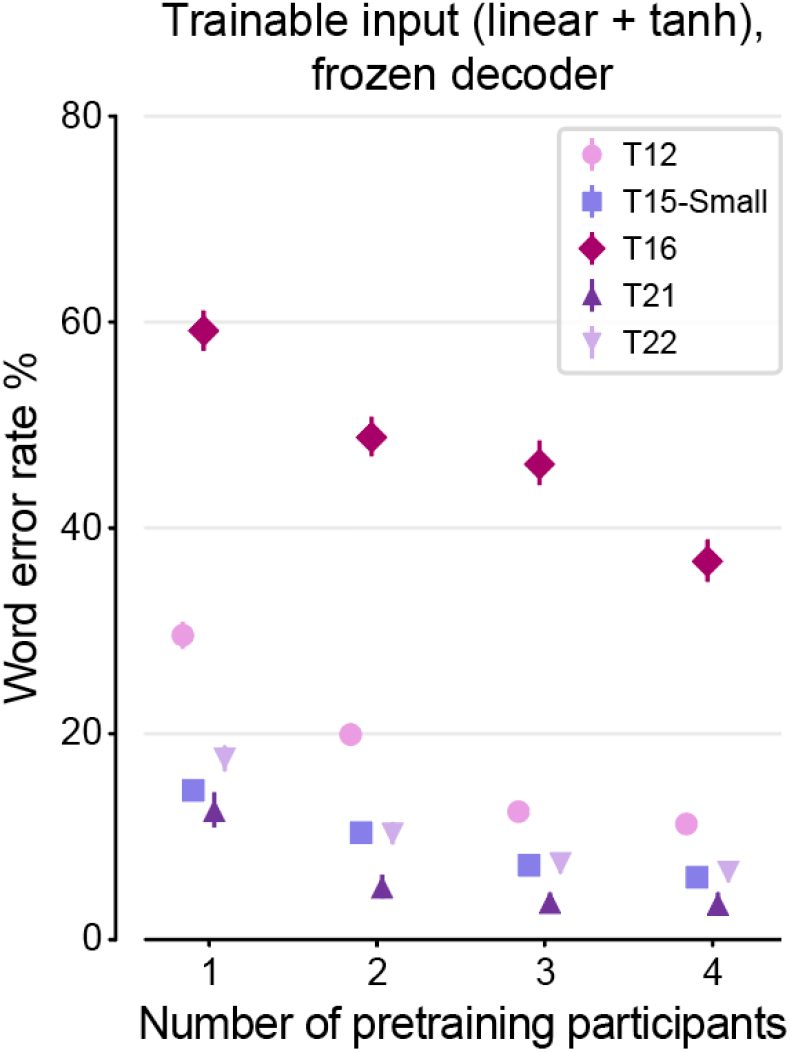
MU model scaling with linear input networks. The same as Fig. 4B, except using simpler input networks (linear projection + tanh nonlinearity) compared to the feed-forward networks used in Fig. 4B.

**Figure S7.**
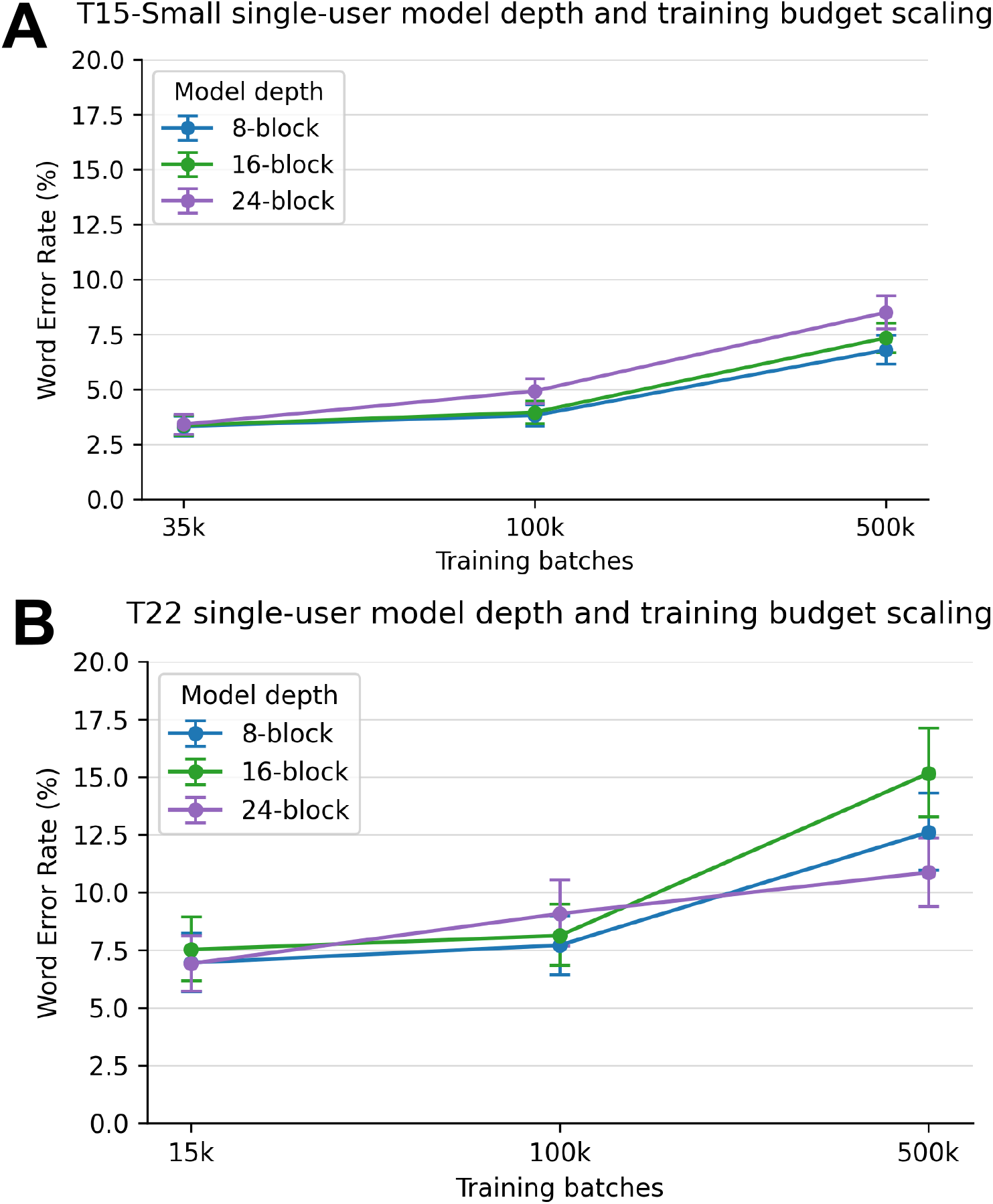
SU model WER as a function of model depth and training budget. SU model WER as a function of model depth and training budget for T15-Small (**A**) and T22 (**B**). Models were trained and tested on the T15-Small and T22 training and test sets described in Table 2. SU models had 8, 16, or 24 decoder blocks and were trained for 35k (for T15) or 15k (for T22), 100k, or 500k training batches. 16 block models used a stochastic depth of 0.3 and had Intermediate-CTC loss computed on decoder blocks 4, 8, and 12. 24 block models used a stochastic depth of 0.3 and had Intermediate-CTC loss computed on decoder blocks 8, 12, 16, and 20. Models trained for 500k batches used maximum AdamW and Muon learning rates of 2e-4 and 6e-4 respectively. All other model architecture and training hyperparameters were identical to those reported for T15-Small and T22 SU models in Table S7. Mean error rates on the held-out test sets are reported with error bars representing 95% confidence intervals.

**Table S1.**
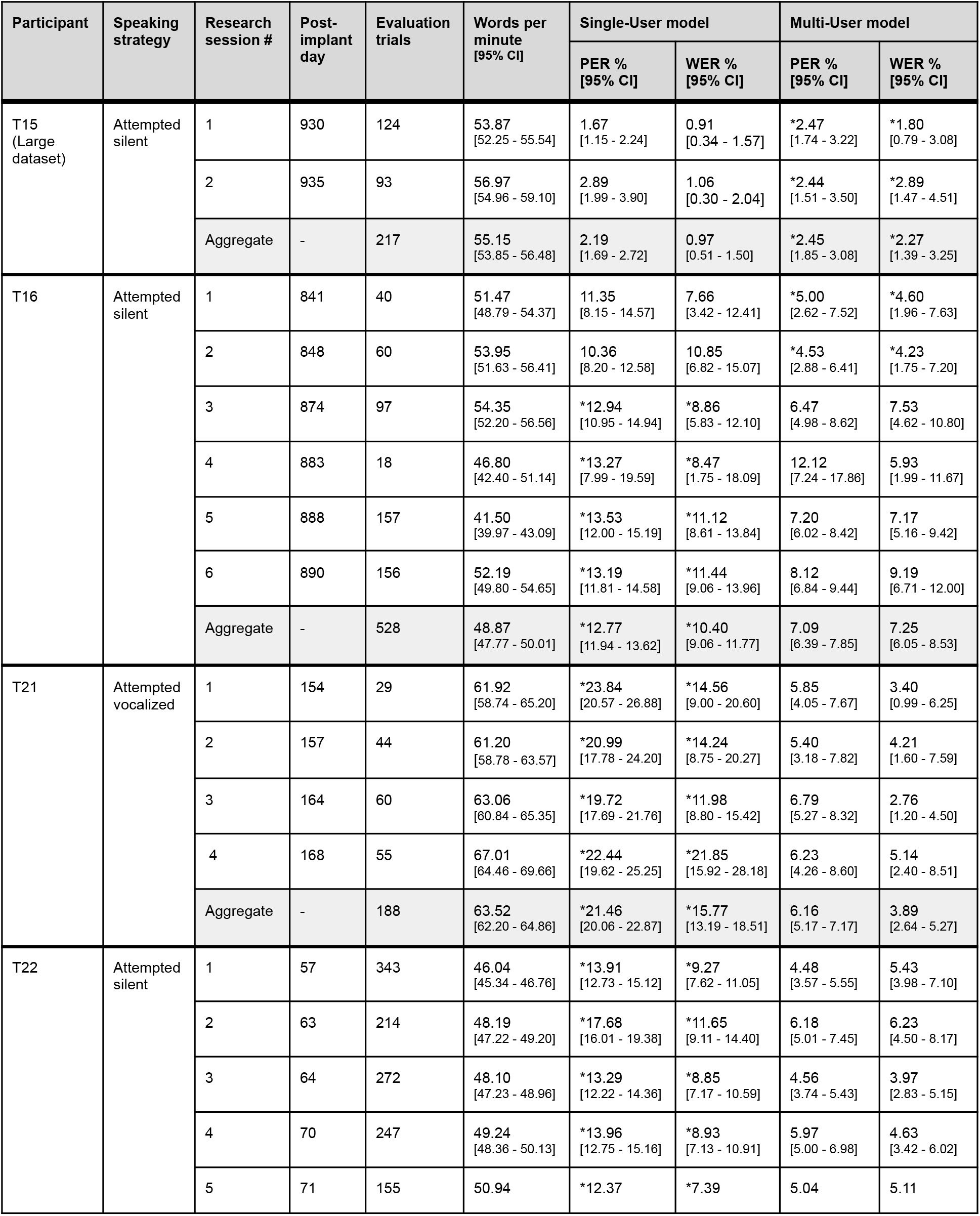
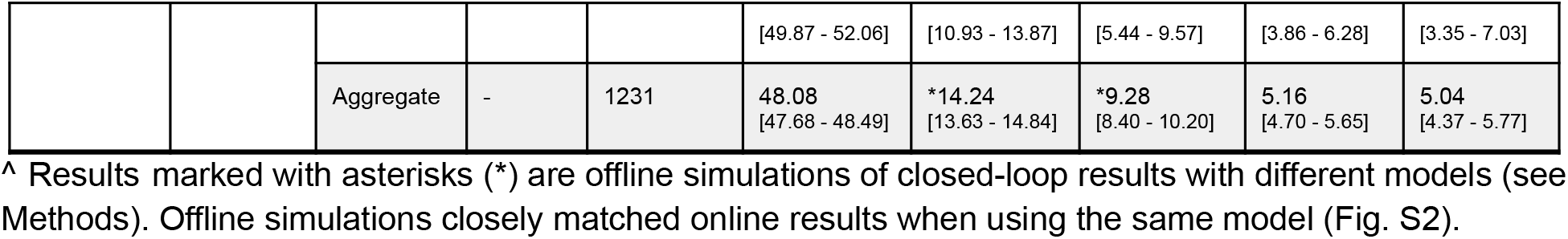
Closed-loop decoding results.

**Table S2.** Closed-loop inference vs. offline re-inference simulation comparison.

| Participant | Evaluation trials | Model Type | Online inference |  | Offline simulation |  |
| --- | --- | --- | --- | --- | --- | --- |
|  |  |  | PER %<br>[95% CI] | WER %<br>[95% CI] | PER %<br>[95% CI] | WER %<br>[95% CI] |
| T15 | 217 | SU | 2.19<br>[1.69 - 2.72] | 0.97<br>[0.51 - 1.50] | 2.31<br>[1.76 - 2.89] | 1.81<br>[1.03 - 2.68] |
| T16 | 100 | SU | 10.76<br>[8.92 - 12.63] | 9.55<br>[6.62 - 12.76] | 10.79<br>[9.29 - 12.32] | 6.10<br>[4.05 - 8.40] |
|  | 428 | MU | 7.66<br>[6.85 - 8.48] | 7.95<br>[6.55 - 9.49] | 5.17<br>[4.48 - 5.90] | 5.40<br>[4.36 - 6.50] |
| T21 | 188 | MU | 6.16<br>[5.17 - 7.17] | 3.89<br>[2.64 - 5.27] | 6.09<br>[5.03 - 7.22] | 4.19<br>[2.96 - 5.52] |
| T22 | 1231 | MU | 5.17<br>[4.70 - 5.65] | 5.04<br>[4.37 - 5.77] | 4.83<br>[4.38 - 5.30] | 5.17<br>[4.47 - 5.91] |

**Table S3.** Offline T21 Personal Use comparison. Evaluating SU and MU models on the same held-out T21 self-generated sentences. The SU model was trained on all T21 sentence data except the last 75% of self-generated trials from three consecutive research sessions (36 sentences total) which were held out as a test set. The MU model was trained on the same data, plus the T12, T15-Large, T16, T17, and T22 training datasets (Table 2). The MU model achieved a mean relative WER improvement of 58.47% compared to the SU model.

| Single-User Model |  | Multi-User Model |  |
| --- | --- | --- | --- |
| PER%<br>[95% CI] | WER%<br>[95% CI] | PER%<br>[95% CI] | WER%<br>[95% CI] |
| 29.25<br>[24.46 - 34.71] | 27.74<br>[18.64 - 38.55] | 11.52<br>[8.06 - 15.57] | 11.61<br>[6.54 - 17.86] |

**Table S4.** Neural phoneme decoder architecture and training procedure ablation. Ablation of the neural phoneme decoder architecture and training procedures introduced since ref. 13, evaluated across single- and multi-user models and multiple participants. Starting from a Base transformer with the architecture and training procedures of ref. 13 (e.g., white-noise augmentation, dropout), we cumulatively add (1) time-masking, time-warping, and channel-masking augmentations; (2) the Intermediate-CTC auxiliary loss and stochastic depth regularization; and (3) weight averaging over the last 66% of validation checkpoints. All training and architecture parameters match Table S7 except those being ablated. Models were trained and tested on the datasets in Table 2, with the MU model trained on T15-Large; to reduce computational cost, the MU model used 12 decoder blocks. MU error rates are averaged across participants. Mean error rates and 95% CIs are computed on seed-averaged test trials across three seeds per model.

| Step | SU T22 |  | SU T15-Small |  | Multi-User Model |  |
| --- | --- | --- | --- | --- | --- | --- |
|  | PER%<br>[95% CI] | WER%<br>[95% CI] | PER%<br>[95% CI] | WER%<br>[95% CI] | PER%<br>[95% CI] | WER%<br>[95% CI] |
| Base transformer | 12.84<br>[11.91 - 13.79] | 13.24<br>[11.61 - 14.92] | 8.33<br>[7.88 - 8.76] | 8.05<br>[7.39 - 8.70] | 9.93<br>[9.54 - 10.33] | 12.32<br>[11.73 - 12.92] |
| + Data Augmentations | 10.11<br>[9.24 - 11.02] | 8.71<br>[7.39 - 10.08] | 7.07<br>[6.64 - 7.50] | 4.70<br>[4.24 - 5.18] | 9.63<br>[9.25 - 10.02] | 10.97<br>[10.40 - 11.55] |
| + Intermediate-CTC & Stochastic Depth | 8.87<br>[8.04 - 9.71] | 7.62<br>[6.45 - 8.83] | 5.67<br>[5.30 - 6.03] | 3.84<br>[3.42 - 4.28] | 6.85<br>[6.52 - 7.19] | 8.37<br>[7.88 - 8.88] |
| + Weight Averaging | 8.54<br>[7.73 - 9.37] | 7.38<br>[6.21 - 8.58] | 5.44<br>[5.08 - 5.80] | 3.33<br>[2.93 - 3.73] | 6.32<br>[6.01 - 6.64] | 7.70<br>[7.24 - 8.17] |

**Table S5.** Training sentence counts after controlling for cross-participant sentence overlap. Training dataset sentence counts before and after controlling for cross-participant sentence overlap. “Train (before)” are the sentence counts of datasets as described in Table 2. “Train (after)” are sentence counts after removing any sentence from any participant’s training dataset that exactly matched any sentence in any participant’s testing dataset.

| Participant | Train (before) | Train (after) | % Decrease |
| --- | --- | --- | --- |
| T12 | 9,680 | 8,040 | 16.94 |
| T15-Large | 278,531 | 259,684 | 6.77 |
| T15-Small | 9,498 | 7,024 | 26.05 |
| T16 | 10,088 | 8,127 | 19.44 |
| T17 | 3,345 | 2,701 | 19.25 |
| T21 | 2,783 | 2,563 | 7.91 |
| T22 | 4,074 | 3,667 | 9.99 |

**Table S6.** Offline decoding comparison when controlling for cross-participant sentence overlap. Offline speech decoding performance comparison between SU and MU models when controlling for cross-participant sentence overlap. We trained SU and MU models with the training datasets described in Table S5. MU models were trained for fewer training steps (300,000 instead of 500,000) and a 12-block model was used instead of a 24-block model to compensate for the lower volume of training data. MU error rates were substantially lower than corresponding SU error rates for all participants with the exception of T15 when compared to a SU model trained on T15-Large. Error rates for all MU and SU models increased when compared to the results reported in Figure 3, likely due to the decreased volume of training data for all participants.

| Participant | Single-User Model |  | Multi-User Model |  |
| --- | --- | --- | --- | --- |
|  | PER%<br>[95% CI] | WER%<br>[95% CI] | PER%<br>[95% CI] | WER%<br>[95% CI] |
| T12 | 12.52<br>[11.94 - 13.11] | 10.19<br>[9.35 - 11.09] | 7.40<br>[6.91 - 7.90] | 7.09<br>[6.44 - 7.79] |
| T15-Large | 1.88<br>[1.68 - 2.09] | 2.64<br>[2.13 - 3.22] | 2.90<br>[2.62 - 3.18] | 3.32<br>[2.76 - 3.93] |
| T15-Small | 8.36<br>[7.88 - 8.84] | 6.24<br>[5.62 - 6.88] | 4.78<br>[4.41 - 5.16] | 3.39<br>[2.95 - 3.85] |
| T16 | 14.77<br>[14.01 - 15.53] | 16.13<br>[14.81 - 17.52] | 11.16<br>[10.45 - 11.89] | 13.89<br>[12.68 - 15.16] |
| T17 | 55.38<br>[54.01 - 56.75] | 75.01<br>[72.37 - 77.61] | 32.23<br>[30.28 - 34.23] | 42.79<br>[39.45 - 46.21] |
| T21 | 17.33<br>[16.10 - 18.61] | 10.71<br>[8.75 - 12.89] | 4.72<br>[3.95 - 5.54] | 3.91<br>[2.79 - 5.17] |
| T22 | 9.60<br>[8.73 - 10.47] | 7.86<br>[6.57 - 9.17] | 4.76<br>[4.04 - 5.51] | 5.47<br>[4.44 - 6.57] |

**Table S7.**
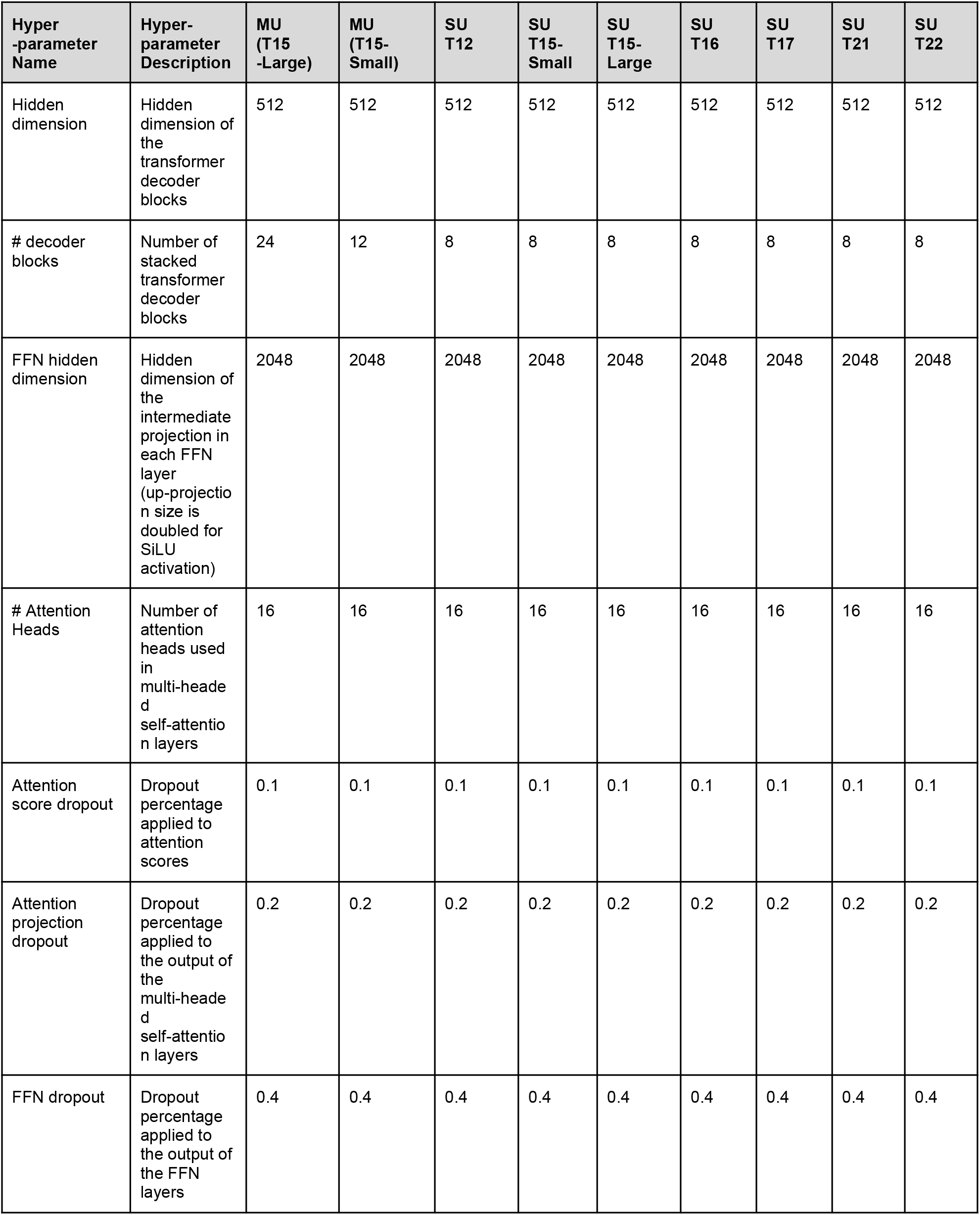
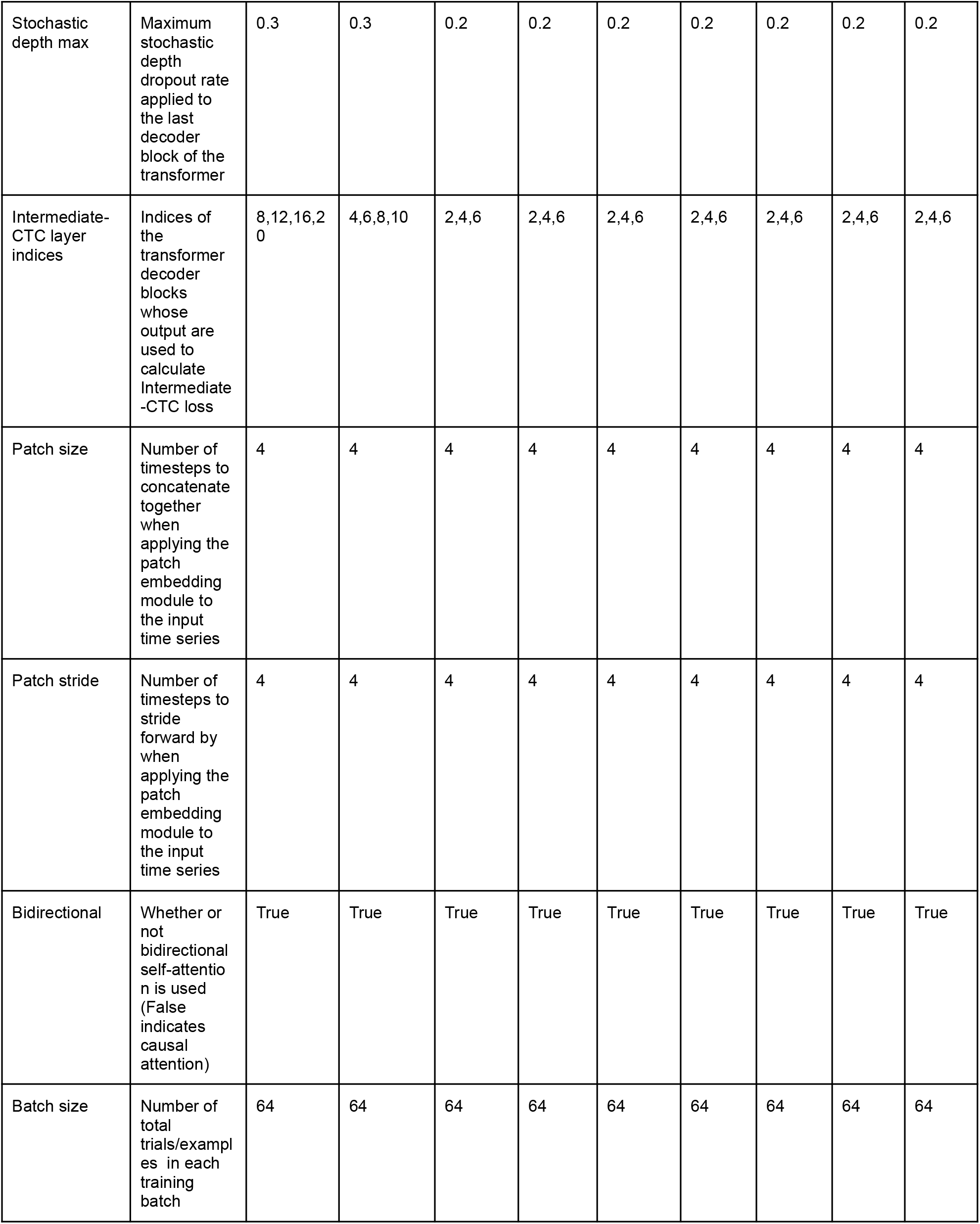
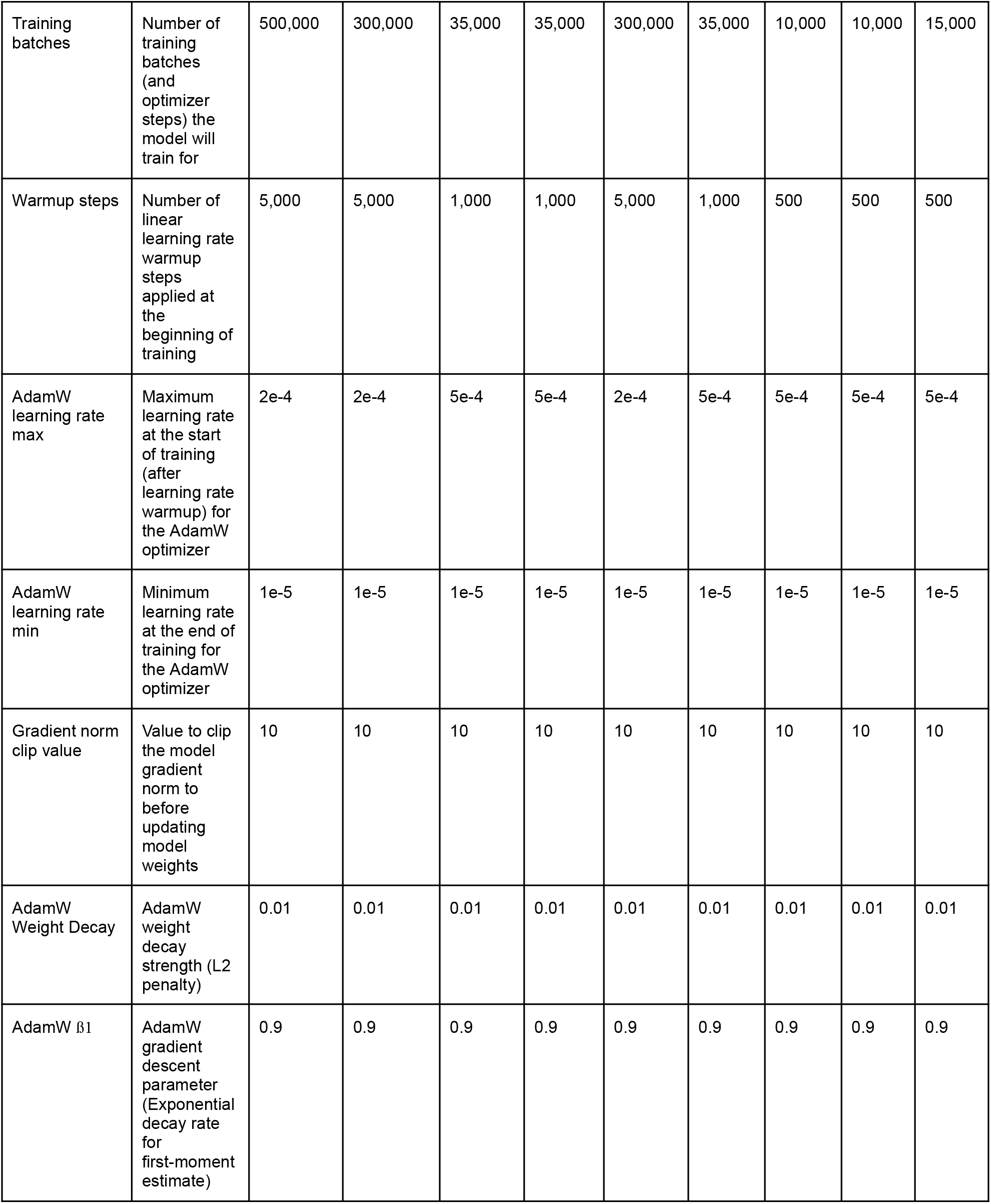
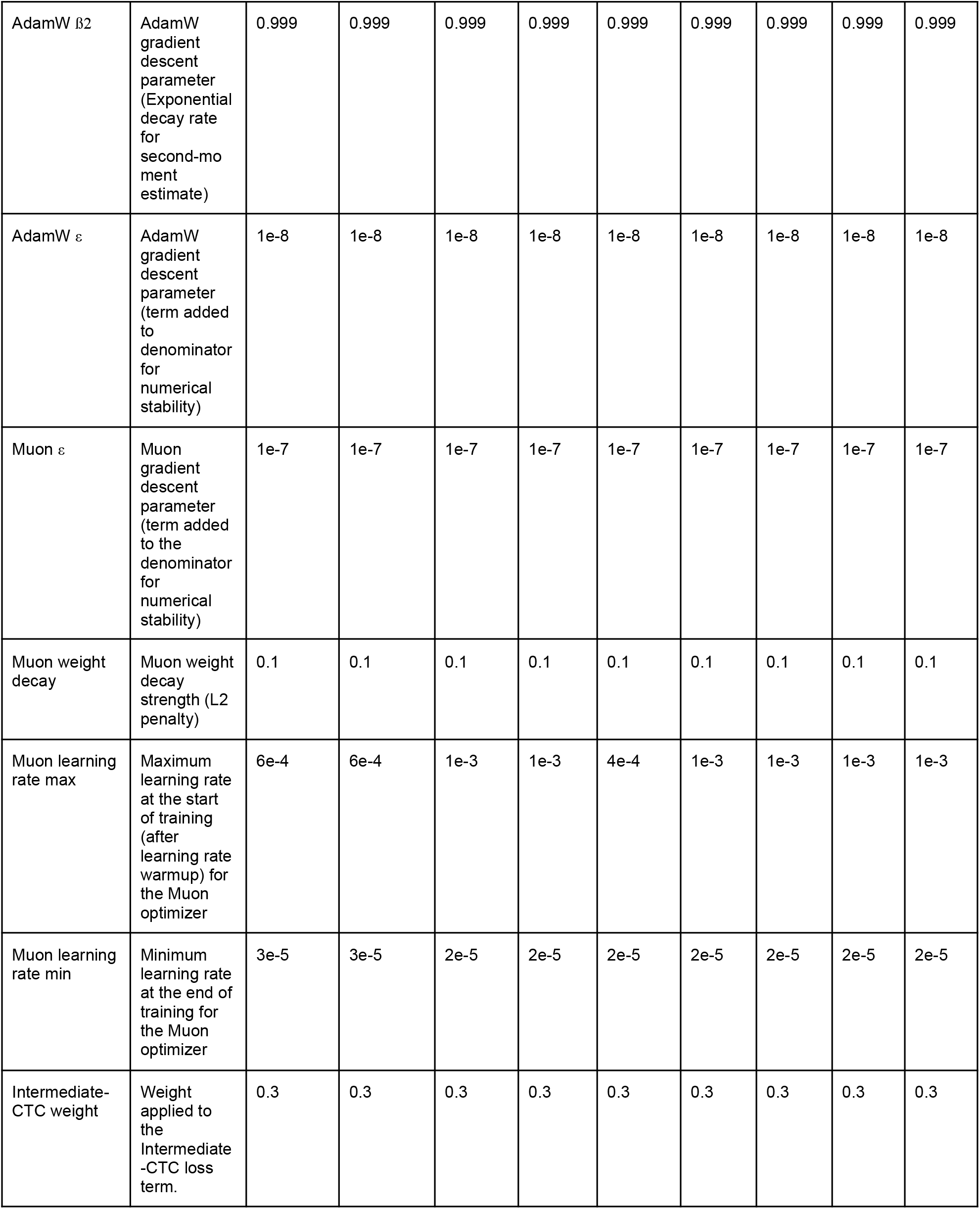
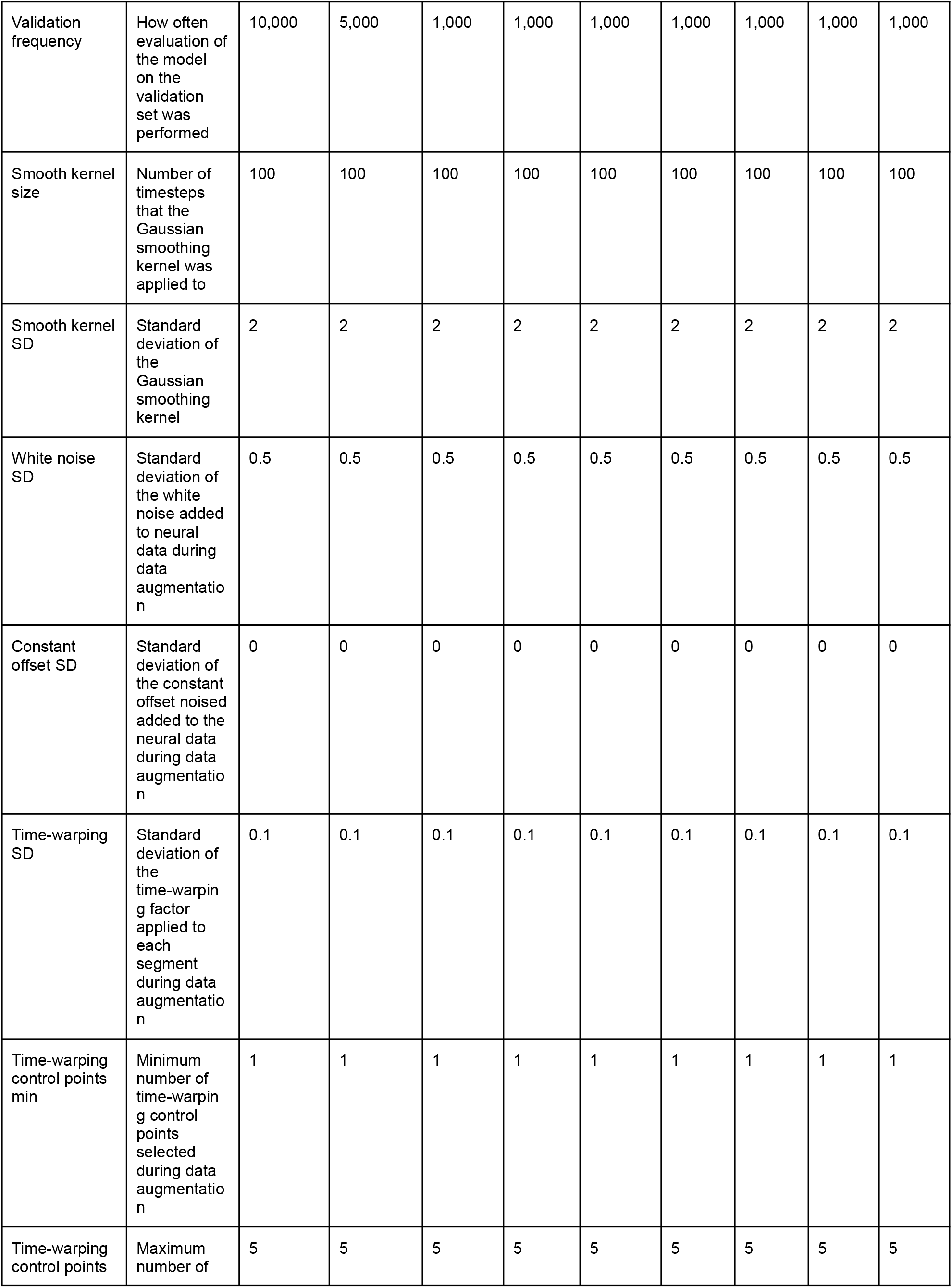
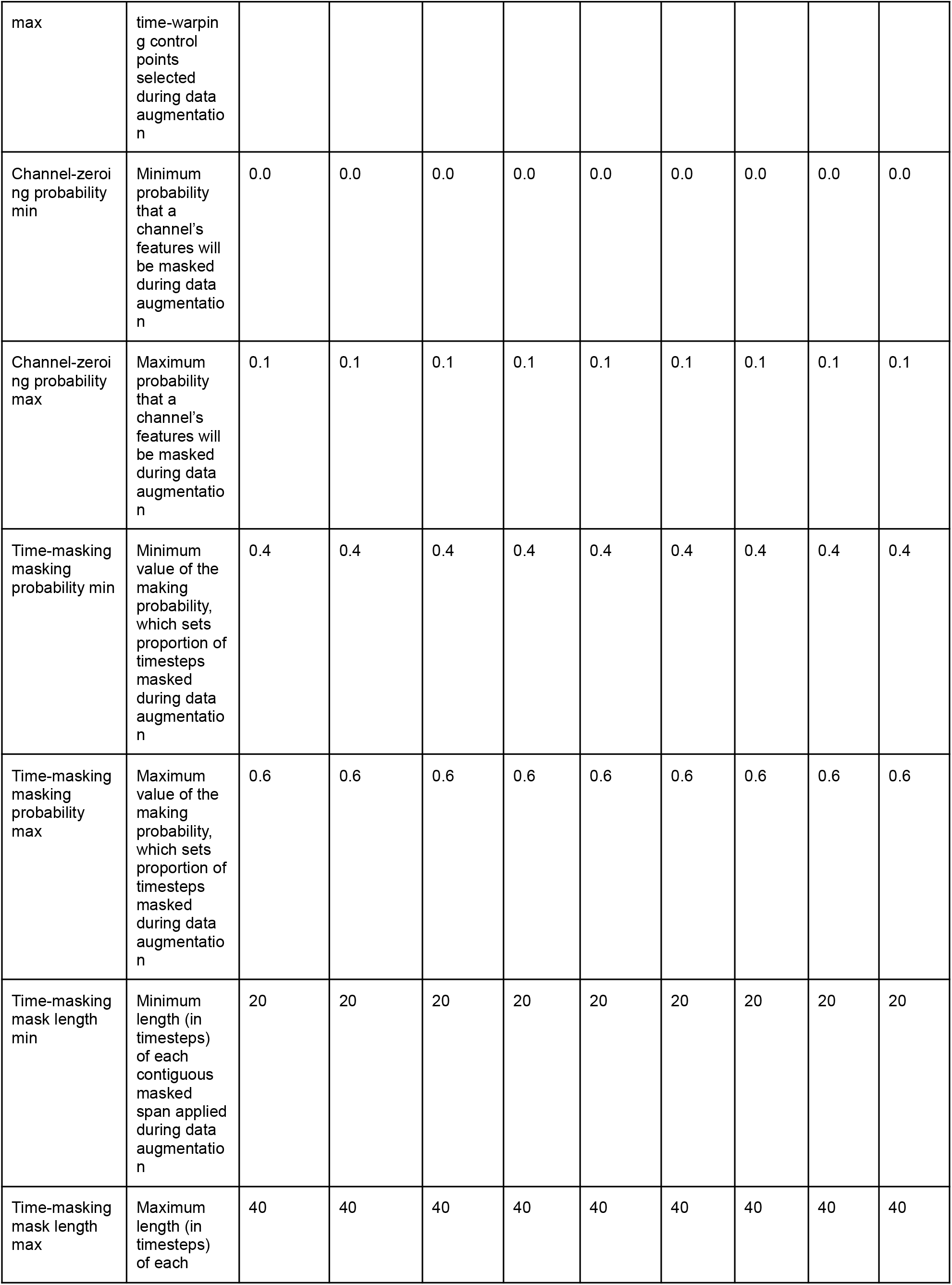
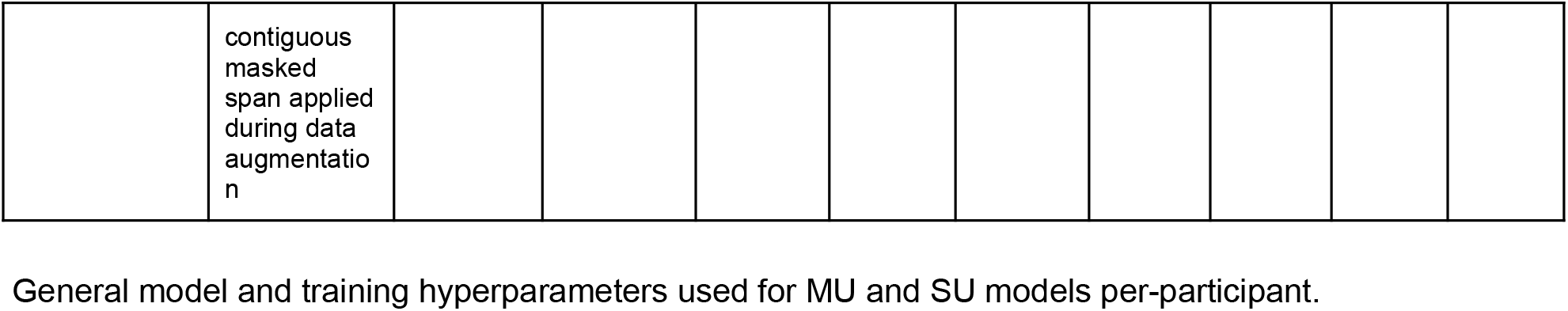
General Multi-user and Single-user model and training hyperparameters

**Table S8.** Single-user model and training hyperparameters for few-shot-learning on held-out participants (. Fig 5.) Model and training hyperparameters for the T21 and T22 SU models used in the few-shot learning analysis (Fig. 5) that differ from those listed for T21 and T22 SU models in Table S7.

| Hyperparameter Name | SU T21 | SU T22 |
| --- | --- | --- |
| # decoder blocks | 4 | 4 |
| FFN hidden dimension | 1024 | 1024 |
| Stochastic depth max | 0.3 | 0.3 |
| Intermediate-CTC layer indices | 2 | 2 |
| Patch size | 5 | 5 |
| Patch stride | 5 | 5 |
| Batch Size | 32 | 32 |
| Training batches | 3,000 | 3,000 |
| Warmup steps | 0 | 0 |
| Muon learning rate max | 1.5e-3 | 1.5e-3 |
| Muon learning rate min | 3e-5 | 3e-5 |
| Validation frequency | 100 | 100 |

**Table S9.** Multi-user frozen-decoder adaptation parameters for few-shot-learning on held-out participants (. Fig 5.) Training hyperparameters for the T21 and T22 MU models used for frozen-decoder adaptation (Fig. 5) that differ from those listed for MU models in Table S7.

| Hyperparameter Name | Value |
| --- | --- |
| Batch Size | 32 |
| Training batches | 1,500 |
| Warmup steps | 0 |
| AdamW learning rate max | 1e-3 |
| Muon learning rate max | 3e-3 |
| Validation frequency | 100 |

**Table S10.**
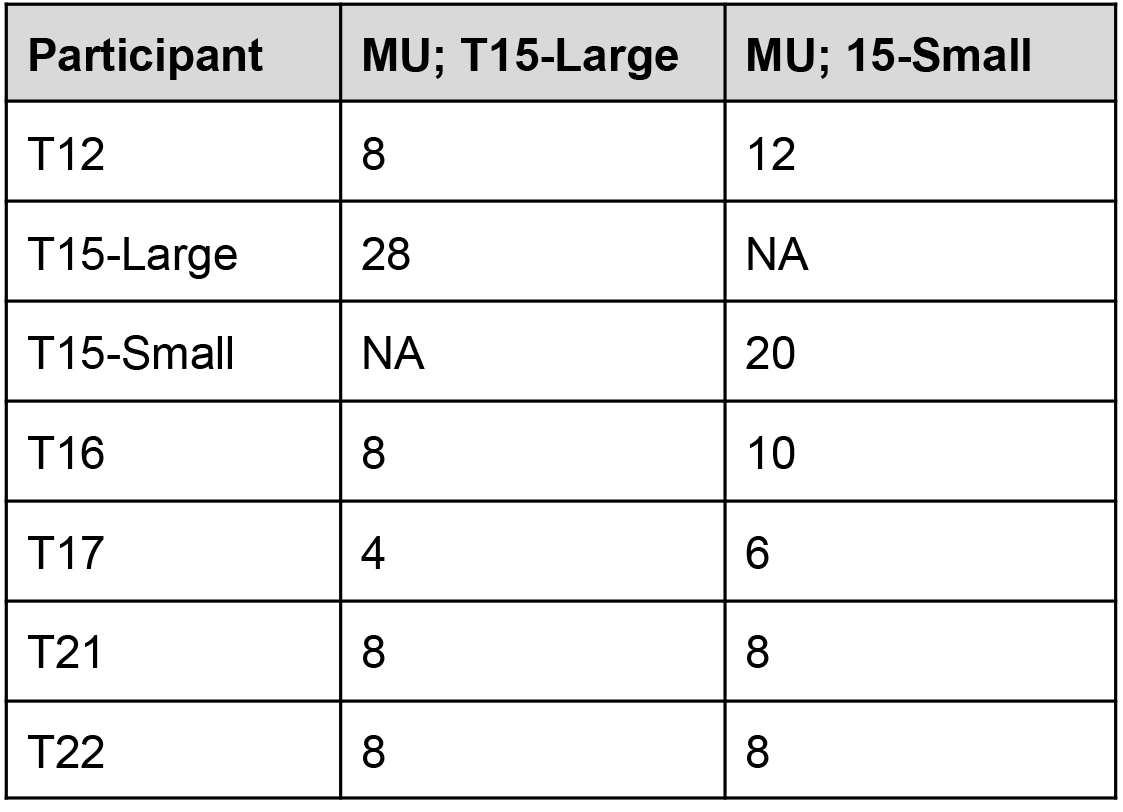
Per-batch sampling rates of participant trials during multi-user training. The number of trials from each participant that were included in each training batch for MU models.

| Participant | MU; T15-Large | MU; 15-Small |
| --- | --- | --- |
| T12 | 8 | 12 |
| T15-Large | 28 | NA |
| T15-Small | NA | 20 |
| T16 | 8 | 10 |
| T17 | 4 | 6 |
| T21 | 8 | 8 |
| T22 | 8 | 8 |

**Table S11.** Offline T21 Right-Hemisphere Only. Comparison of speech decoding performance between a SU model and a finetuned MU model on T21’s two right-hemispheric arrays. The MU model was pretrained on T12, T15-Large, T16, and T22 and then finetuned via trainable-decoder adaptation only on data from T21’s two right-hemispheric arrays. The SU model was trained from scratch only on data from T21’s two right-hemispheric arrays. Besides the exclusion of neural features from T21’s four left-hemispheric arrays, training and test sets are the same as specified in Table 2. Mean error rates and 95% CIs are computed on seed-averaged test trials across three seeds per model.

| Single-User Model |  | Multi-User Model |  |
| --- | --- | --- | --- |
| PER%<br>[95% CI] | WER%<br>[95% CI] | PER%<br>[95% CI] | WER%<br>[95% CI] |
| 50.97<br>[49.76 - 52.20] | 68.08<br>[65.51 - 70.64] | 39.37<br>[37.72 - 41.05] | 47.97<br>[44.82 - 51.30] |

**Table S12.**
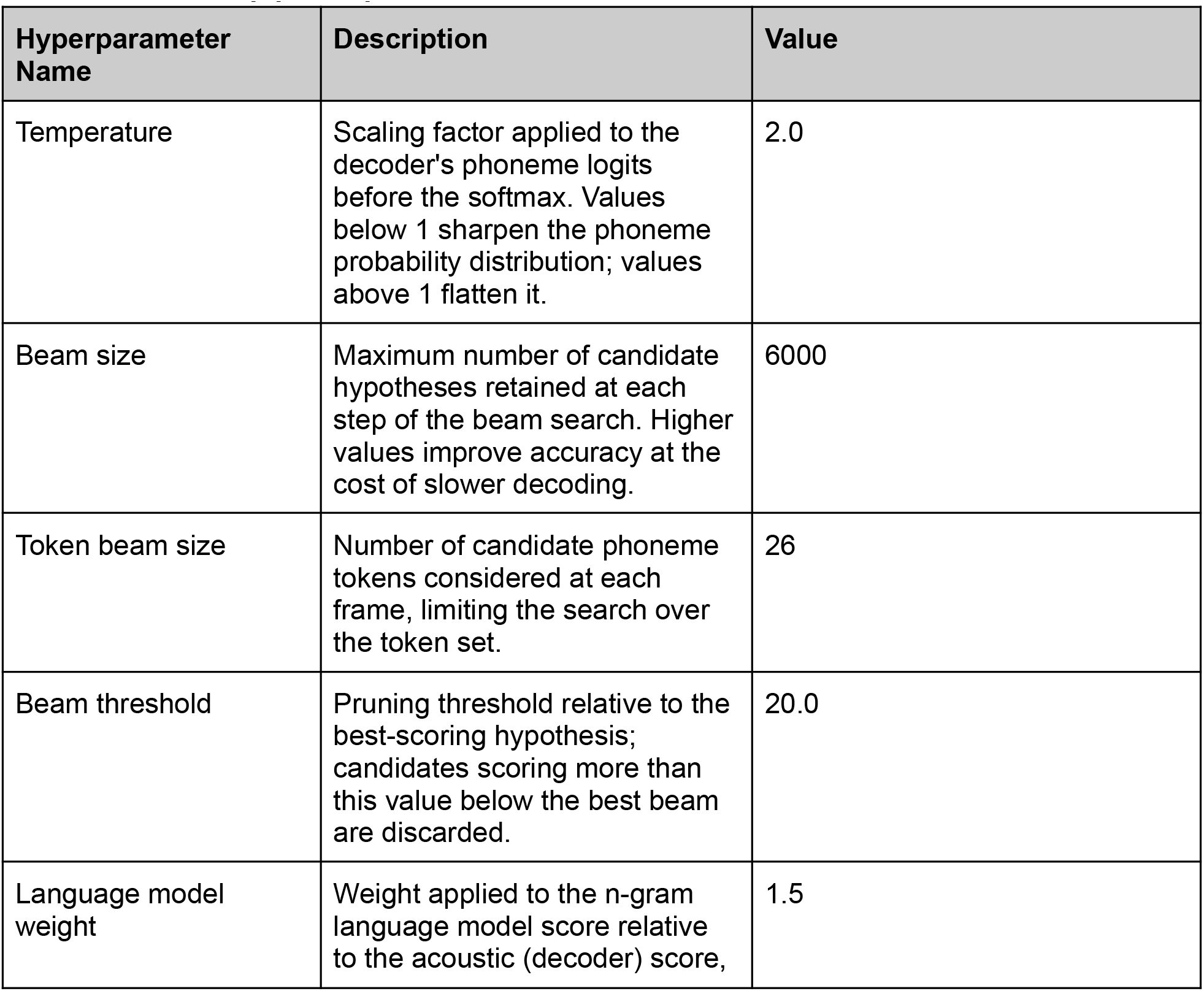
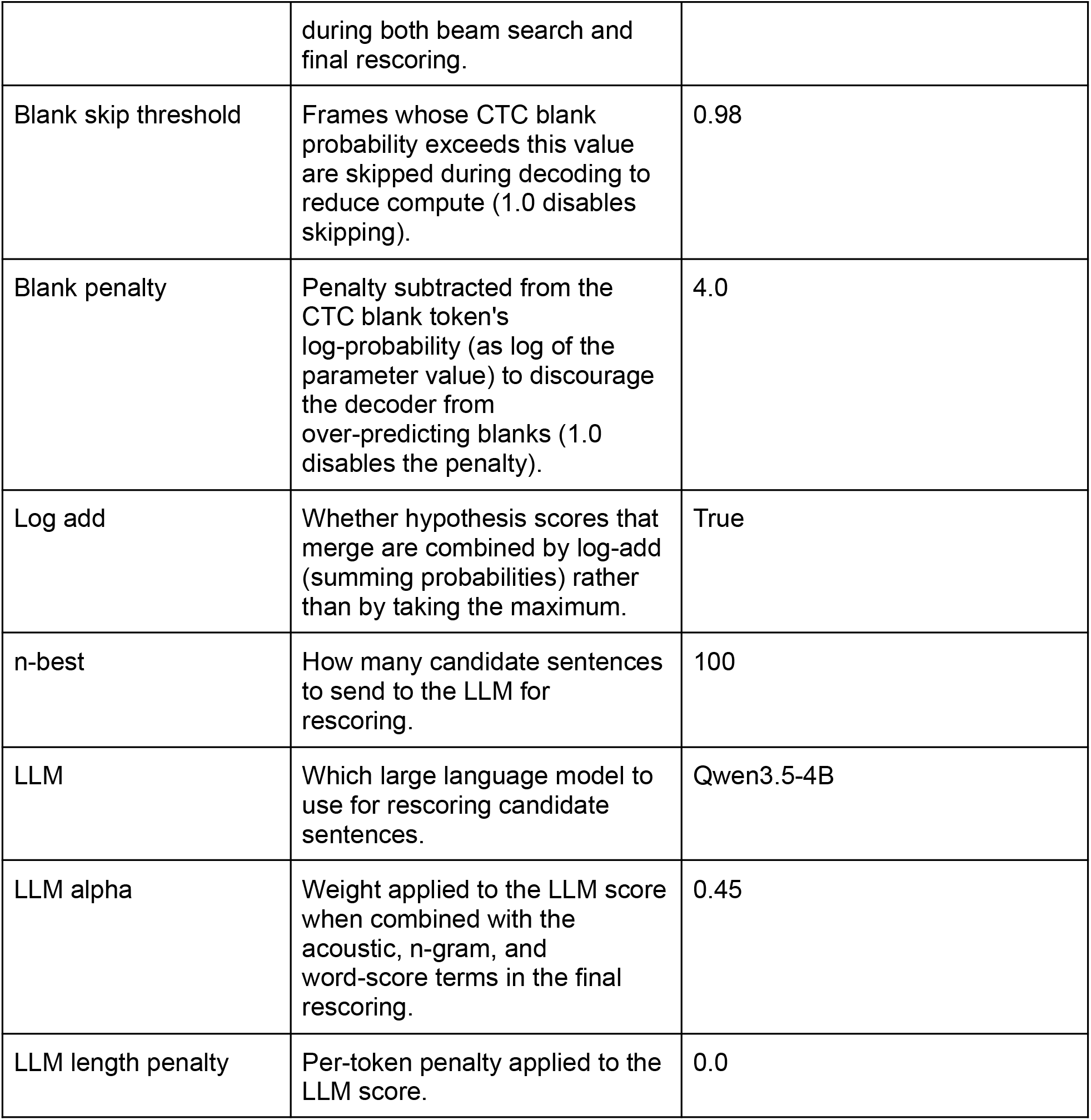
KenLM pipeline parameters.

